# AURKB-driven dissolution of CIZ1-RNA assemblies from the inactive X chromosome in mitosis

**DOI:** 10.1101/2025.09.17.676856

**Authors:** Lewis Byrom, Gabrielle L. Turvey, Adam A. Dowle, Megan Thomas, Navin Shirodkar, Ben J. Green, Maxwell Brown, Charlotte Ball, Kate E. Chapman, Elena Guglielmi, William Dickson, Emma Noon, Sajad Sofi, Justin F-X. Ainscough, Alfred A. Antson, Dawn Coverley

## Abstract

Cip1-interacting zinc-finger protein 1 (CIZ1) interacts with *Xist* lncRNA to form large RNA-protein assemblies at the inactive X-chromosome (Xi) in female mammalian nuclei, plus smaller assemblies in both sexes. CIZ1 assemblies influence underlying chromatin, and their disruption alters the expression of autosomal and X-linked gene clusters. Here, we explore the regulated dissolution of CIZ1-Xi assemblies during mitosis and show that, like *Xist*, CIZ1 is released in prometaphase under the regulation of Aurora Kinase B (AURKB). The part of human/mouse CIZ1 comprising 179/181 C-terminal amino-acids encodes a matrin-3 domain that facilitates dimerization to form a compact folded core with disordered C-terminal extensions. Mass spectrometry revealed 56 high-confidence interacting partners of the C-terminal fragment, predominantly chromatin, nuclear matrix and RNA-binding proteins. Phosphomimetic mutation of three conserved AURKB sites in the C-terminal extensions released CIZ1 from its nuclear anchor points, but did not affect its interaction with chromatin or nuclear matrix proteins. In contrast, the same mutations, or deletion of the C-terminal extensions, abolished interaction with RNAs including *Xist*. Together, the data suggest CIZ1 is a regulatable component of the protein-RNA assemblies that preserve epigenetic stability across the nucleus, and that AURKB drives their dissolution in mitosis via dissociation of CIZ1 from RNA.

**Bullets:** The data show regulated dissolution of RNA-protein assemblies involved in protection of epigenetic state.

RNA spatially constrains CIZ1 assemblies via multivalent interactions at sub-nuclear sites.

CIZ1 dimerizes to present dual extensions whose phosphorylation by AURKB dissolves RNA interaction in mitosis.

**Graphical abstract:** 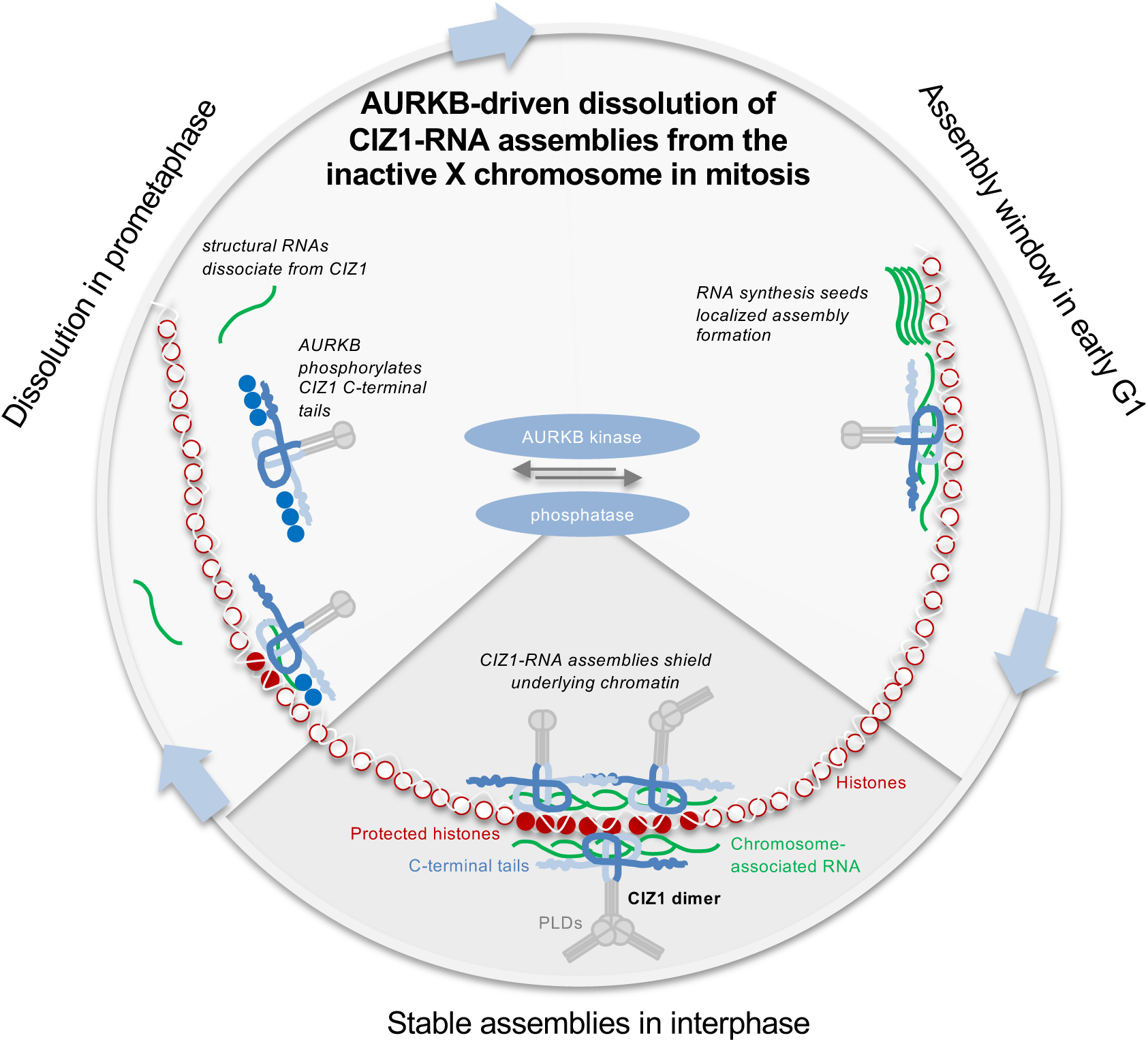

## Introduction

The mammalian cell nucleus is organised to facilitate tight control over gene expression - some genes are accessible, some inaccessible and some poised to respond to changing cues. Control is achieved through associated proteins and their post-translational modifications, but also via non-coding RNAs that seed the formation of protein assemblies that are an integral part of the nuclear architecture, and crucial determinants of structure-based control. Historically, the proteins whose location within the nucleus remain unaffected by extraction of DNA and lipids were referred to as nuclear matrix proteins, and those that are sensitive to digestion of RNA, the RNA-dependent nuclear matrix (1,2). Many of these are part of RNA-seeded protein condensates, and include those that assemble around the archetypal long non-coding RNA (lncRNA) *Xist* at the inactive X chromosome (Xi) in female mammalian cells (3). RNA-protein particles (RNPs) have been implicated in all aspects of nuclear function, including RNA modification, processing, turnover and localisation, as well as in the control of gene expression. While most RNPs do not diffuse freely, there is no consensus on the extent to which they remain spatially isolated verses connected into a wider network. Fundamental questions remain around how a cell, in which chromatin is spatially and structurally organised according to its lineage, can collapse its nucleus and condense chromosomes at mitosis, and then reassemble accurate structural organisation in daughter cells.

Our focus is Cip1-interacting zinc finger protein 1 (CIZ1), which meets the definition of a nuclear matrix protein in that it remains *in situ* in sub-nuclear foci even when chromatin is digested away (4). CIZ1 sub-nuclear foci are evident in both sexes, but in females they also aggregate into supramolecular assemblies around the Xi, seeded by *Xist* lncRNA. In differentiated cells, in which *Xist*-dependent repression of Xi chromatin has been achieved, CIZ1 and *Xist* co-exist within RNP particles, and their recruitment to Xi is co-dependent (5,6). Although CIZ1 is one of the most stable components of the multiprotein supramolecular assemblies (SMACs) that drive heterochromatinization of the Xi (7), neither the loss of *Xist* nor deletion of CIZ1 causes widespread de-repression of X-linked genes once its repressed state has been established, yet both are required for accurate control over a subset of ‘escape’ genes. In the case of CIZ1 these are not limited to X-linked genes but encompass approximately 0.4% of the transcriptome in primary embryonic fibroblasts (PEFs) (8) distributed across all chromosomes, suggesting that the assemblies that are associated with autosomes perform a similar function to those at the Xi, likely as part of assemblies seeded by other lncRNAs.

CIZ1 interacts directly with *Xist*, preferring its E-repeat sequences (5,6), but also interacts with other RNAs via at least two RNA interaction interfaces (9). In the N-terminal half of CIZ1, RNA interaction is mediated by low-complexity prion-like domains that also drive self-association, and in the C-terminal third by as yet undefined sequences. Both are required for *de novo* formation of CIZ1 assemblies at Xi in fibroblasts, and in the co-recruitment of *Xist* (9) (Fig.1A). The requirement for multivalent interaction with RNA makes CIZ1-RNA assemblies susceptible to the effect of monovalent competitors, and assemblies can in fact be dissolved experimentally using dominant-negative fragments of CIZ1 (DNFs), with immediate (within days) effects on underlying chromatin and gene expression (10). Unscheduled dissolution of CIZ1-RNA assemblies is now linked with early-stage breast cancers (10), so the molecular determinants of their normal assembly and disassembly within the nucleus is important to understand. Here, we explore the interactions that support anchorage, and the regulated dissolution of CIZ1-Xi assemblies during mitosis in normal cells.

**Fig. 1.**
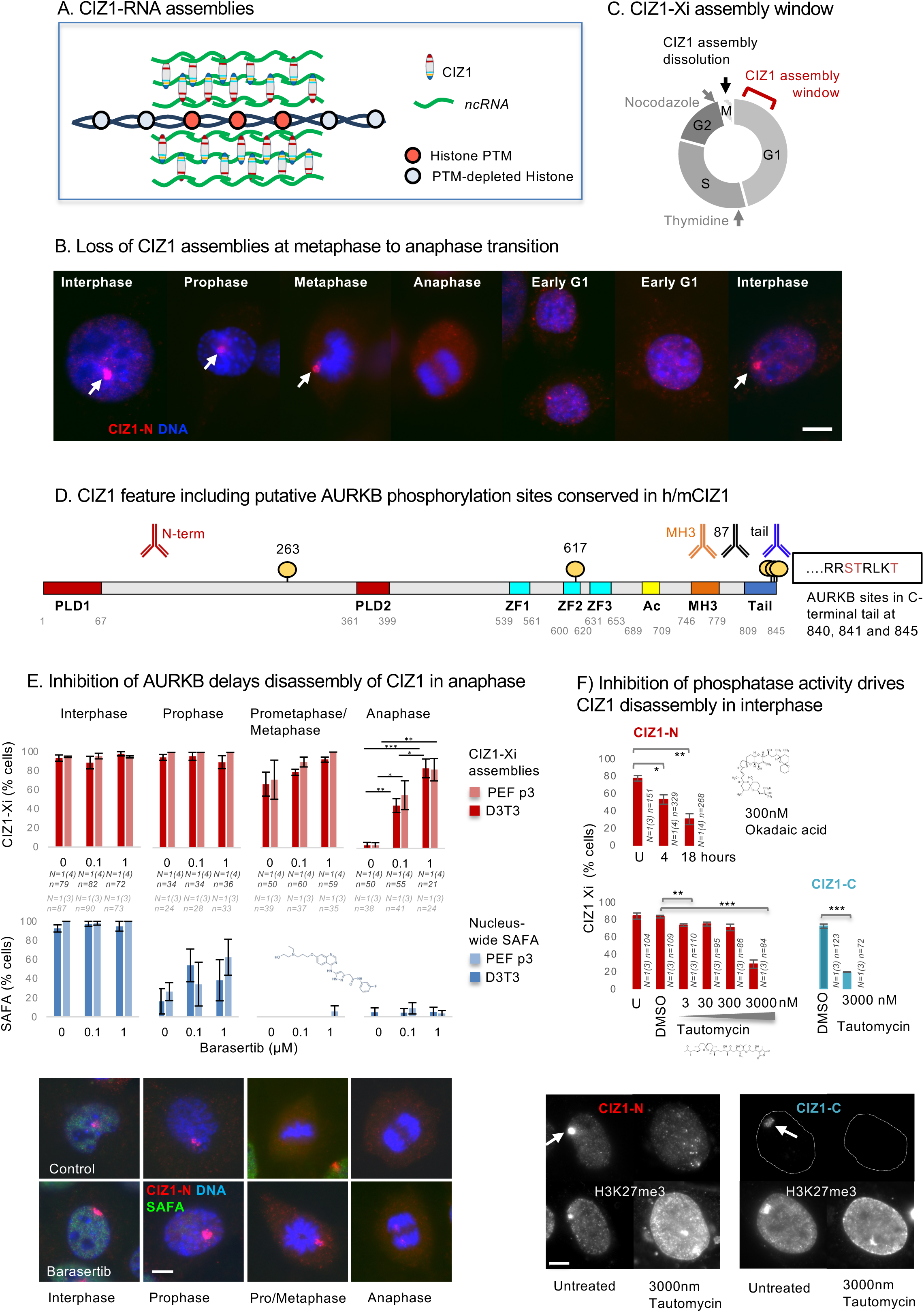
Dissolution of CIZ1-Xi assemblies in mitosis. A) Model CIZ1-RNA assemblies surrounding and protecting the modification status of underlying chromatin (9,10). B) Illustrative immunofluorescence images of female murine D3T3 cells stained for CIZ1 via N-terminal epitopes (CIZ1-N, red, detected with pAb 1794), revealing large protein assemblies at the inactive-X chromosome (white arrows), that are not detected in mitosis. DNA is blue, bar is 5µm. C) Diagram illustrating loss in mitosis and the early G1 phase window during which reformation of CIZ1-Xi assemblies takes place (10). D) Map showing conserved putative AURKB phosphorylation sites between murine and human CIZ1 (circles) displayed on full-length murine CIZ1 (NP_082688.1.) The location of epitopes of CIZ1 antibodies used throughout are shown above. Conserved prion-like domains (PLD1 and PLD2) are in red (9), Zinc fingers 1-3 in cyan (ZnF_C2H2 SM00355, ZF_C2H2 sd00020 and ZF_C2H2 sd00020), acidic region (Ac) in yellow, Matrin-3 homology domain (MH3) in orange (ZnF_U1, smart00451), and h37/m38 amino-acid C-terminal tail in blue. The sequence context and identity of three conserved AURKB phosphorylation sites in the extreme C-terminus are shown. E) Frequency of cells with CIZ1-Xi assemblies (red), or nucleus-wide SAFA (blue), in cells passing through the stages of mitosis indicated, for D3T3 cells and female primary embryonic fibroblasts (PEFs at p3), in the presence and absence of the AURKB kinase inhibitor barasertib (30) at 0.1 and 1uM. Results show the average of 3-4 independent replicates within one experiment for each line, with SEM. n indicates the number of nuclei inspected (PEF grey, 3T3 black). Statistical analysis of CIZ1-Xi frequency in anaphase cells shows one-way ANOVA with Tukey post hoc test within each cell type, where * <0.05, ** <0.01, *** <0.001. Below, example immunofluorescence images of D3T3 cells through mitosis, with and without 1uM barasertib. Cells were stained for the N-terminal domains of CIZ1 (CIZ1-N, red) and SAFA (green). DNA is blue, bar is 5 microns. F) Upper histogram shows the proportion of cells with CIZ1-marked Xi in cycling populations of female D3T3 cells after the indicated times exposed to 300nM Okadaic acid, visualized via CIZ1-N (red). Lower, histograms show the effect of the indicated concentrations of Tautomycin for 15 hours, stained for CIZ1-N or the ‘tail’ epitope in the C-terminal end of CIZ1 (CIZ1-C, rabbit pAb). Comparison of technical replicates is by T-test, where * <0.05, ** <0.01, *** <0.001. Error bars show SEM. Below, example images showing H3K27me3-marked Xi chromatin in cells stained for CIZ1-N or CIZ1-C, at 15 hours with or without tautomycin. Bar is 5 microns.

## Methods

### Recombinant protein expression and purification

N-terminal GST-tagged CIZ1 constructs were expressed in transformed BL21 RP Codon plus cells (Agilent). Cells were grown in Luria Broth starter cultures supplemented with 100 µg/mL ampicillin at 37°C for 16 hours whilst shaking at 200 RPM. Starter cultures were used to inoculate 1L expression cultures grown until OD600 = 0.10 – 0.15. Expression was initiated by lactose driven autoinduction for 24 hr at 20°C whilst shaking at 200 RPM. Cultures were harvested via centrifugation at 4500 RPM, 4°C for 15 min (Thermo Scientific, Fiberlite F9-6 x 1000 LEX Fixed Angle Rotor, 096-061075). The resulting bacterial pellet was gently washed in PBS wash buffer (PBS supplemented with 1 mM DTT, 3 mM EDTA, 1 mM PMSF). Washed cell pellets were centrifuged at 4500 RPM (Thermo Scientific, Fiberlite F15-8 x 50cy Fixed Angle Rotor, 75003663) 4°C for 15 min, the supernatant was discarded, and the pellet snap frozen in liquid nitrogen until purification. Cells were thawed on ice and resuspended in PBS wash buffer supplemented with EZblock protease inhibitor, at a ratio of 5 mL buffer per 1g E. coli. Cells were sonicated using a 6 mm diameter MS 73 microtip (Bandelin) and a SONOPULS ultrasonic homogeniser (Bandelin) operating at 60-70 % power. Sonication (30 seconds on then 30 seconds off) was repeated five times. Sonicated cell lysates were centrifuged at 15000 RPM (Thermo Scientific, F15-8 rotor) 4°C for 30 min. The supernatant was removed then syringe filtered using a 0.45 µm filter. Protein was purified via GST affinity chromatography using ӒKTA pure™ (GE healthcare) chromatography systems at room temperature with chilled buffers, or in a 4°C cold cabinet. 5 mL GSTrap 4B column (Cytiva) were equilibrated with 10 column volumes (cv) PBS wash buffer before applying the clarified supernatant at 0.5 mL / min. The column was subsequently washed with 10 column volumes (cv) PBS wash buffer followed by 10 cv cleavage buffer (50 mM Tris-HCl pH 7, 150 mM NaCl, 1mM EDTA, 1 mM DTT). The column was removed and directly injected with 1 cv cleavage buffer supplemented with 0.4% v/v PreScission Protease (Cytiva) for on-column GST-tag cleavage at 4 °C for 16 hours. Untagged CIZ1 protein was eluted on AKTA with 2 cv. Eluted protein was concentrated to ∼ 0.5 mL using Vivaspin® 6, 10 kDa MWCO columns and directly injected via a 0.5 or 1 mL loop onto a pre-equilibrated Superdex^®^ 200 Increase 10/300 GL column (Cytiva). Protein was separated over 1 cv at 0.5 mL/min, collecting 0.5 or 1 mL elution fractions. Peak fractions were identified using absorbance at 280 nm, and were pooled and concentrated before snap freezing in liquid nitrogen for storage at -80 °C. NanoDrop spectrophotometer readings were used to measure final protein concentration and purity was verified by SDS-PAGE.

To prepare bait protein for interaction studies, expression of GST-tagged C-terminal CIZ1 fragments or GST controls were carried out as described above, with resuspension of cells in cleavage buffer supplemented with protease inhibitor and 1 mM PMSF. Clarified supernatants (∼40 mL) were incubated with Glutathione Sepharose 4B bead slurry (Cytiva) at 4°C for 1 hour rotating at 7 rpm, in the presence of 0.8 µL Benzonase (Millipore) and 8 µL RNAse (Roche). GST-CIZ1 coated beads were extensively washed in cleavage buffer followed by isotonic buffer before addition of nuclear extracts.

### SEC-MALLS

For size-exclusion chromatography multi-angle laser light scattering, 100 μL of >1mg/mL purified protein was analysed using an Superdex® S200 10/300 GL column (GE Healthcare), equilibrated in cleavage buffer with a flow rate of 0.5 mL/min using a Shimadzu HPLC system (SPD-20A UV detector, LC20-AD isocratic pump system, DGU-20A3 degasser and SIL-20A autosampler) at room temperature (20 ±2°C). Detection of light scattering was carried out using a Wyatt HELEOS-II multi-angle light scattering detector and a Wyatt rEX refractive index detector. Solvent was 0.2 μm filtered before use and a further 0.1 μm filter was present in the flow path. The column was equilibrated with at least 2 column volumes of solvent before use and flow was continued at the working flow rate until baselines for UV, light scattering and refractive index detectors were all stable. Shimadzu LabSolutions software was used to control the HPLC and Astra 7 software for the HELEOS-II and rEX detectors. The Astra data collection was 1 minute shorter than the LC solutions run to maintain synchronisation. MWs were estimated using the Zimm fit method with degree 1, using the Astra 7 software. A value of 0.182 was used for protein refractive index increment (dn/dc).

### Cells, culture and transfection

Primary Embryonic Fibroblasts (PEFs), D3T3 murine fibroblast cell line (11) and HeLa cells (12) (Table 1) were cultured as adherent populations in High Glucose DMEM GlutaMAX (Gibco), supplemented with 10% FBS (PAA) and 1% Pen/Strep/Glutamine (Gibco), at 37°C, 5% CO2. Cells were passaged before reaching 90% confluency using D-PBS (Gibco) and trypsin (Gibco). PEFs were generated from C57BL/6 mice embryos at day 13-14 of gestation. CIZ1 null mice were generated from ES clone (IST13830B6, TIGM) using a Neomycin resistance gene trap downstream of exon 1, as described previously (5). Heterozygous CIZ1+/- mice were bred to generate CIZ1+/+(Wild-type) mice and CIZ1-/- (CIZ1 null mice) and CIZ1 status confirmed for parents and embryo-derived primary cells by a combination of PCR genotyping, immunoblot and immunofluorescence to detect CIZ1 protein. Cells utilised for transfection and immunofluorescence assays were plated onto glass coverslips and used at approximately 80% confluence. Transfection reactions were carried out in DMEM supplemented with 11% Opti-MEM (Gibco), 0.3% TransIT-X2 (Mirus), for 24 or 48 hrs as indicated. Typically, 200 ng pEGFP-CIZ1 plasmid or derived fragment was used per coverslip in a 0.5ml volume.

**Table 1.** Cells.

| Cell type | Strain ID | CIZ1 Genotype | Sex |
| --- | --- | --- | --- |
|  | **Cell Lines** |  |  |
| Murine fibroblast | D3T3 |  | Female |
| Human carcinoma | HeLa |  | Female |
|  | **Primary cell isolates** |  |  |
| PEF | 13.49 | WT | Female |
| PEF | 13.47 | WT | Male |
| PEF | 35.2A | KO | Female |
| PEF | 13.66 | KO | Female |
| PEF | 13.54 | KO | Male |
| PEF | 13.85 | KO | Male |
| PEF | 13.79 | WT | Female |
| PEF | 13.67 | KO | Female |
| PEF | 13.55 | KO | Female |
| PEF | 13.37 | WT | Female |
| PEF | 45.1fb | WT | Male |
| PEF | 16.5de | KO | Male |
| PEF | 13.74 | KO | Female |
| PEF | 13.54 | KO | Male |
| PEF | 13.84 | WT | Male |

### Nuclear extracts

S3 HeLa cells at 90% confluency were washed in PBS then rinsed in cold isotonic buffer (50 mM Hepes pH 7.8, 5 mM MgCl_2_, 5 mM potassium acetate, 135 mM NaCl), followed by cold isotonic buffer with protease inhibitor (Roche) for 10 mins at 4°C. Cells in minimal buffer were scrape harvested at 4°C and disrupted by dounce homogenisation in isotonic buffer, until nuclei were released. Protein fractions were extracted via centrifugation to separate insoluble (nuclei and cytoskeleton), and soluble fractions. The insoluble fraction was resuspended in a cell-equivalent volume of isotonic buffer supplemented with 400 mM NaCl for 10 mins on ice, then centrifuged to generate a supernatent fraction. The pellet was resuspended in a half cell-equivalent volume 800 mM NaCl then centrifuged to generate a second supernatent fraction. Soluble protein fractions collected in 400 and 800 mM NaCl were pooled, and diluted to a final concentration of 135 mM NaCl before incubation with GST-CIZ1 beads for 2 hr at 4°C rotating at 7 rpm. Beads were washed five times in isotonic buffer before snap freezing. Three independent studies compared GST-C179_WT_ to a C179-derived mutant, and were each controlled by comparison to proteins retrieved by GST alone.

### Immunofluorescence assays

Cells grown on coverslips were washed in D-PBS followed by a brief wash with cytoskeletal buffer (CSK, 10 mM PIPES pH 6.8, 100 mM NaCl, 300 mM Sucrose, 1 mM sucrose, 1 mM MgCl2, 1 mM EGTA), or CSKD (CSK supplemented with 0.1% triton X100). Cells were fixed with 4% w/v paraformaldehyde (PFA) for 15 min followed by two D-PBS washes. For analysis of mitotic cells coverslips were fixed in PFA without prior wash steps. Following fixation, cells were washed in D-PBS and then incubated in BSA antibody buffer (PBS supplemented with 0.1% BSA, 0.02 % SDS, 0.1% Triton X100) for 30 min. Primary antibodies (Table 3) were diluted in BSA antibody buffer and incubated with cells on coverslips for 1 hr at 37°C, washed three times with BSA antibody buffer, followed by secondary antibodies for 1 hr at 37°C, then washed with BSA antibody buffer followed by D-PBS. Coverslips were mounted onto glass slides with DAPI-containing Vectashield (Vector Laboratories).

### Microscopy

All images were taken using a 63X/1.40 plan-apochromat objective and Zeiss filter sets 2, 10, 15 (G365 FT395 LP420, BP450-490 FT510 BP-515-565, BP546/12 FT580 LP590). Image acquisition was carried out using an Axiocam 506 mono camera (Zeiss) and Axiovision software (SE64 version 4.9.1). Within experiments where fluorescence intensities were quantified, exposures were kept consistent across a set of conditions. Fluorescence microscopy images were analysed and prepared for figures using FIJI and ImageJ2 version 2.14.0/1.54f, or ImageJ version 1.53. Quantification of fluorescence from immunostained cells used unedited raw images. The DAPI channel was utilised to create an image mask to extract both nuclear area measurements, and a region within which fluorescence intensity mean, minimum and maxima were extracted.

Live cell imaging to quantify morphological differences between wildtype and CIZ1-null primary cell populations over the cell cycle was conducted using LiveCyte 2 (PhaseFocus). PEFs at passage 1 were cultured in high glucose DMEM then split into 24-well glass bottom plates (Cellvis) a day prior to imaging. Culture media was replenished immediately prior to imaging of two wells per cell population across three fields of view each (1000 µm), using a 10X Plan N objective. Exposure time was set at 25 ms at 100% power, with slice count of 1 and slice spacing of 5.0 µm. Images were taken at 6 min intervals over 24 hours. Visual estimates of the number of cells entering mitosis and the number of cells successfully dividing over a time window were made by identifying cells that brighten and round up. Differences between wildtype and CIZ1-null populations were quantified by inspecting before and after this point in time. Entry into mitosis was determined by counting the total number of cells in the first frame and then calculating the total number of mitotic cells observed over the timeframe as a percentage. Successful division was determined based on whether mitotic cells flattened out to produced two daughters, generating three categories: normal (a clean division into two equally sized daughters), abnormal (an aberrant division into two irregular or asymmetric daughters), and failed (flattening of mitotic cells without division). Significant differences between the wildtype and CIZ1-null cell populations were evaluated by paired t-test.

### RNA probe generation

*Xist* digoxygenin (DIG)-labelled RNA probes were synthesised by generating cDNA for *Xist* repeat E via PCR from pCMV-Xist-PA (Addgene) (13), using primers in Table 2 to incorporate a T7 promoter sequence. GADPH RNA probe was generated from pTRI-GAPDH-Mouse control plasmid (Invitrogen Northern Max kit) and Human 18S Ribosomal RNA probe was generated from Transcription Kit control plasmid, all described previously (9). PCR products (3-4 μg) or control plasmid were used directly for *in vitro* transcription with MEGAshortscriptTM Transcription Kit (Ambion). 20 µL reactions contained 7.5 mM ATP, 7.5 mM CTP, 7.5 mM GTP, 7.5 mM UTP, 0.5 mM dig-11-UTP (Roche) made to 1x transcription buffer in nuclease free water, including 2 μL T7 Megashortscript enzyme mixand were incubated at 37°C for 4 hr. 2 U TURBO DNase was added to completed reactions to remove DNA template for 15-30 min at 37°C. DIG labelled RNA probes were purified from transcription reactions using MEGAclear Transcription Clean-Up Kit (Ambion). RNA was collected in 2x 40 μL elutions, using pre-heated (95°C) buffer and quantified by measuring absorbance at 260 nm.

**Table 2.**
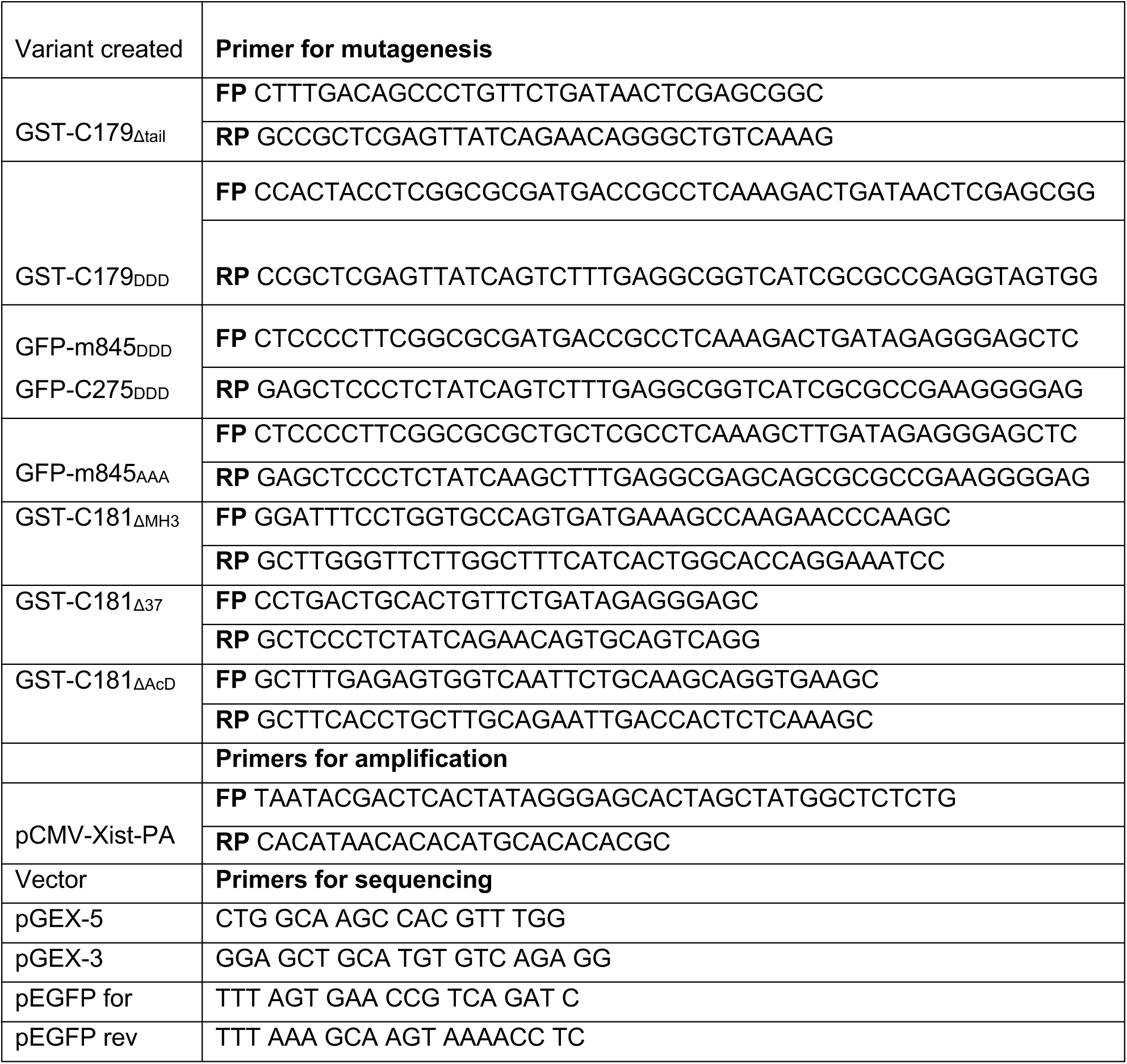
Primers.

### Electrophoretic mobility shift assays (EMSAs)

Purified proteins in binding buffer (10 mM Tris-HCl pH 7 at 25°C, 30 mM NaCl, 2.5 mM MgCl2, 0.1 % IGEPAL CA-630, 0.1 mg/mL yeast tRNA, 1.5 % RNase OUT (Invitrogen), 0.2 mM EDTA and 0.2 mM DTT) were heated to 30 °C for 20 min. RNA probes were diluted and denatured at 80 °C for 3 min before snap cooling on ice and used at final concentrations in reactions at 0.66 nM for Xist repeat E, 0.65 nM for GAPDH and 0.63 nM for 18S rRNA. RNA was incubated with proteins for 20 min at 30 °C, then loading dye was added before application to a 0.8 % w/v agarose gel prepared in TBE running buffer, for electrophoresis at 75V for 65 min at 4°C. Gels were transferred to nylon membrane (Hybond-N+) via weighted capillary action for 45 mins, then following incubation with 2x SSC buffer (SLS) were membranes were crosslinked via UV (125 mJ for 80 sec). Membranes were washed for 2-5 mins in wash buffer, followed by blocking buffer for 30-35 mins (Roche). Anti-dig Fab fragment (Roche) was diluted 1:10,000 (v/v) in blocking buffer and incubated with the membrane for 30 mins then washed twice for 15 mins before equilibration in detection buffer for 5 mins. RNA mobility was revealed by incubation with 0.25 mM chemiluminescent substrate chloro-5-bysubstituted adamantly-1,2-dioxetane phosphate substrate (CSPD Roche) for 10 min at 37 °C, and visualized on a PXi gel imaging system (Syngene). RNA probe quantification was carried out using FIJI, analysing the raw integrated density values of captured images, and expressed as percentage using the control lane of 0 μM protein as reference,

### Western blots

Protein samples were denatured in SDS-PAGE loading buffer at 90°C for 5-10 mins then separated through Mini-PROTEAN TGX precast gels 4-15% (BIORAD), using SDS-PAGE running buffer (25 mM Tris, 192 mM Glycine, 0.1% SDS) at 100V for 1-2 hrs. Whole cell lysates were separated at 40V for 20 mins then 60V for 1-2 hrs. PageRuler Plus Prestained Protein Ladder (Thermo scientific) was used throughout to monitor migration. Separated proteins were transferred to nitrocellulose (GE healthcare) using iBlot2 transfer system (Invitrogen). Blots were blocked in antibody appropriate buffers; 5% BSA, 5% or 10% non-fat dried milk diluted in phosphate or Tris buffered saline (PBS or TBS) with 0.1% Tween20. Blots were incubated with diluted primary antibodies (Table 3) overnight at 4°C or 1-2 hr at RT, then washed in blocking buffer three times before incubation for 1 hr with HRP-conjugated secondary antibodies. Blots were incubated with ECL chemiluminescence substrate (Thermo Scientific) and signal revealed using PXi gel imaging system (Syngene) with Genesys V1.8.2.0 software. Quantification of western blots was carried out using Gene Tools software 4.3.14.0 (Syngene). For visualisation of protein content gels were stained with Coomassie blue (Invitrogen), and membranes with ponceau S stain (Merck).

**Table 3.** Antibodies.

| Target | Source |
| --- | --- |
| CIZ1 (C-terminal tail) Rabbit anti-peptide | Novus (NB100-74624) |
| CIZ1 (N-term, 1794) Rabbit anti-recombinant CIZ1 | (11) |
| CIZ1 (C-terminal tail) Goat anti-peptide | this study |
| CIZ1 (C-terminus, 87) Mouse anti-recombinant CIZ1 | (9) |
| CIZ1 (MH3 domain) | Biorbyt (orb524815) |
| SAFA | Abcam (ab10297) |
| SAFA | Proteintech (14599-1-AP) |
| SAFB2 | Proteintech (11642-1-AP) |
| SAFB2 | Santa Cruz Biotechnology (sc-514963) |
| H2AK119ub1 | Cell Signalling Technology (D27C4) |
| Tri-Methyl-Histone H3 (Lys27) | Cell Signalling Technology (C36B11) |
| α-Tubulin | Abcam (ab7291) |
| Aurora Kinase B (AURKB) | Novus (NBP2-50039) |
| SMARCA5 | Biorbyt orb340809 |
| NUMA1 | Antibodies.com (A12645) |
| Histone H3 | Abcam (Ab1791) |
| GFP | Sigma (SAB4301138) |
| Actin | Abcam (ab11003) |
| Actin | Sigma (A4700-2mL) |
| GST | Abcam (ab9085) |
| Goat anti-rabbit IgG Alexa Fluor 568 (Red) | Invitrogen (A-11011) |
| Goat anti-mouse IgG Alexa Fluor 488 (Green) | Invitrogen (A-11001) |
| Goat anti-rabbit IgG Alexa Fluor 488 | Invitrogen (A11034) |
| Goat anti-mouse IgG Alexa Fluor 568 | Invitrogen (A11031) |
| Donkey anti-rabbit IgG Alexa Fluor 568 (Red) | Invitrogen (A-10042) |
| Donkey anti-goat IgG Alexa Fluor 488 (Green) | Invitrogen (A-11055) |
| HRP-IgG anti rabbit | Jackson Immuno (211-032-171) |
| HRP-IgG anti mouse | Jackson Immuno (155-035-174) |

**Table 4.**
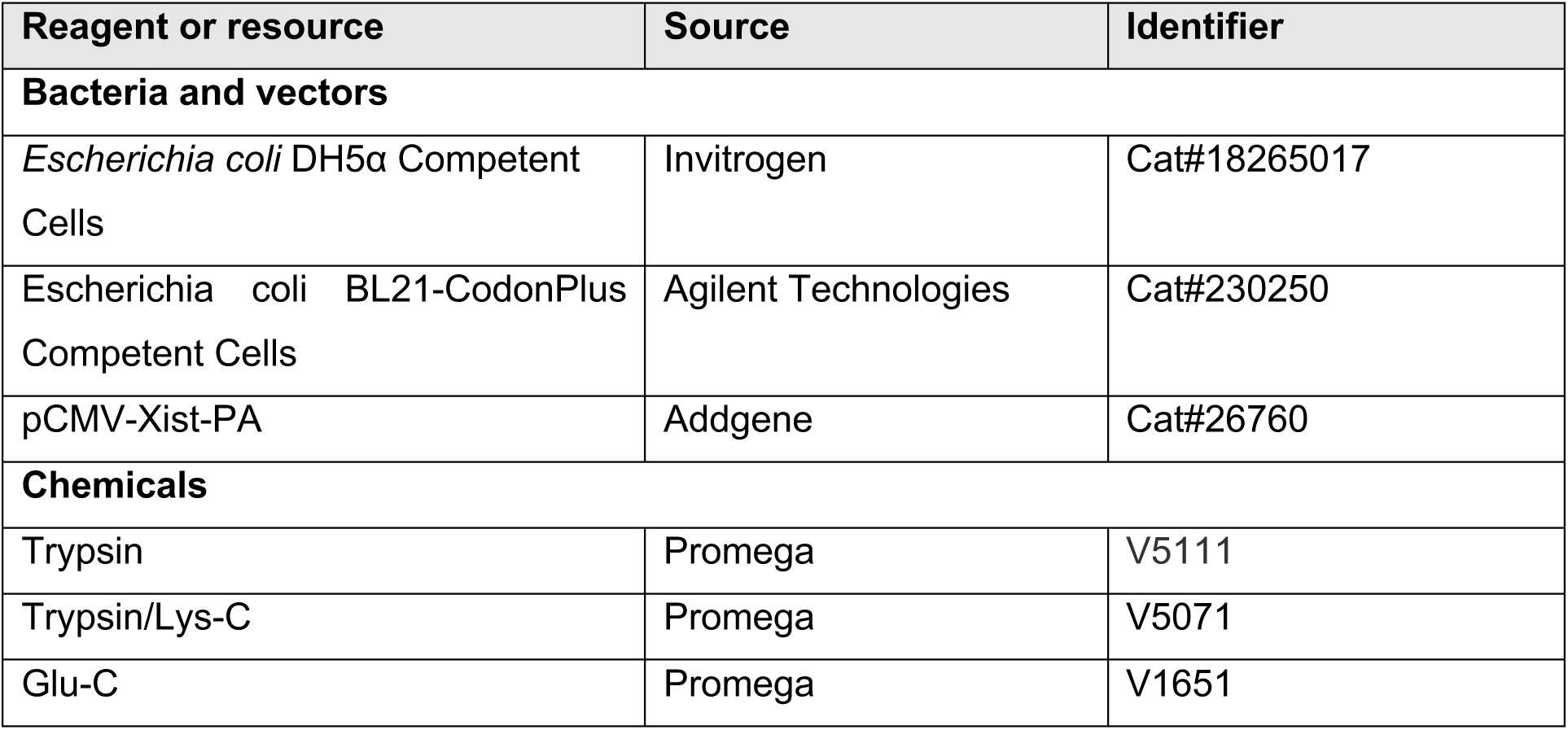
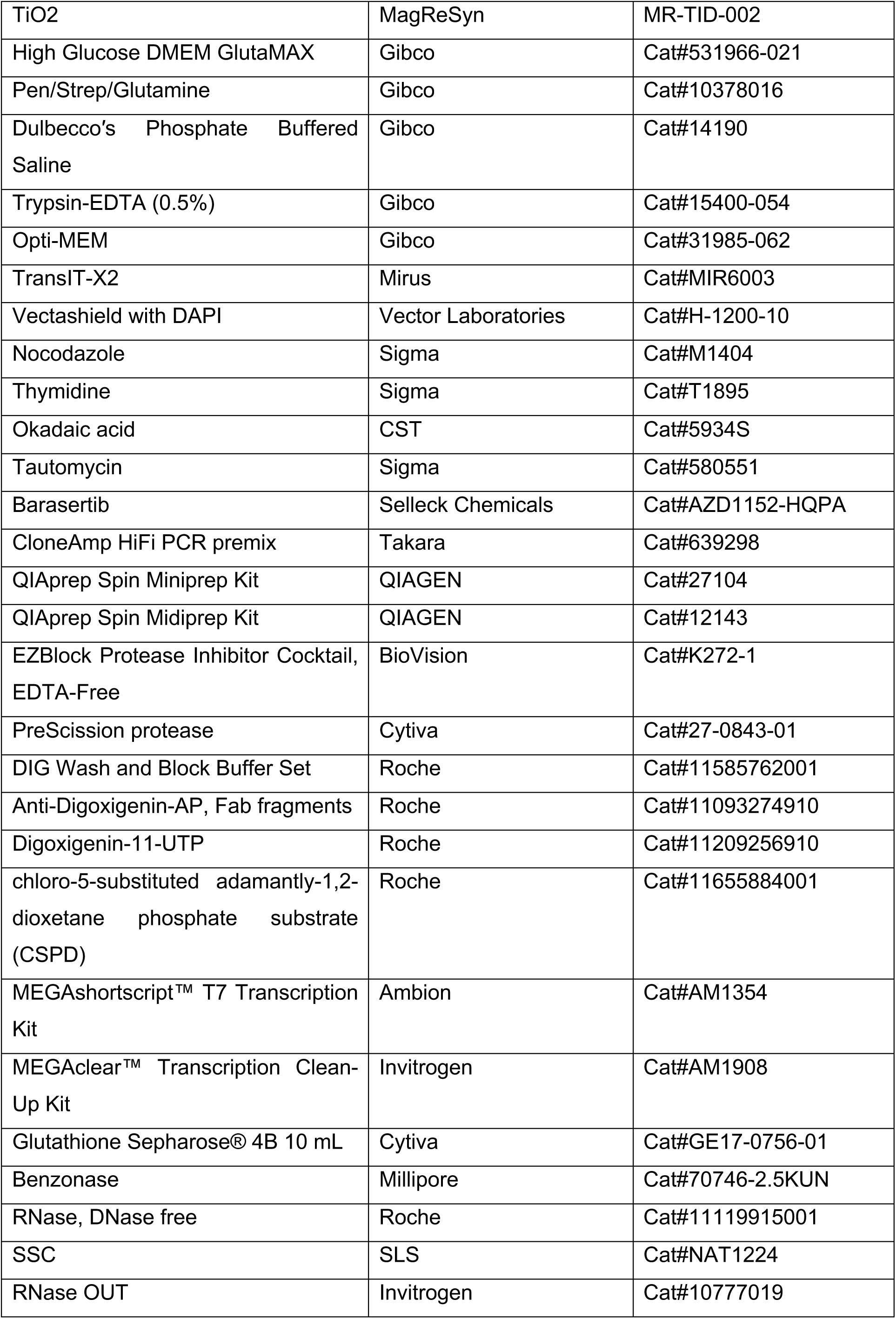
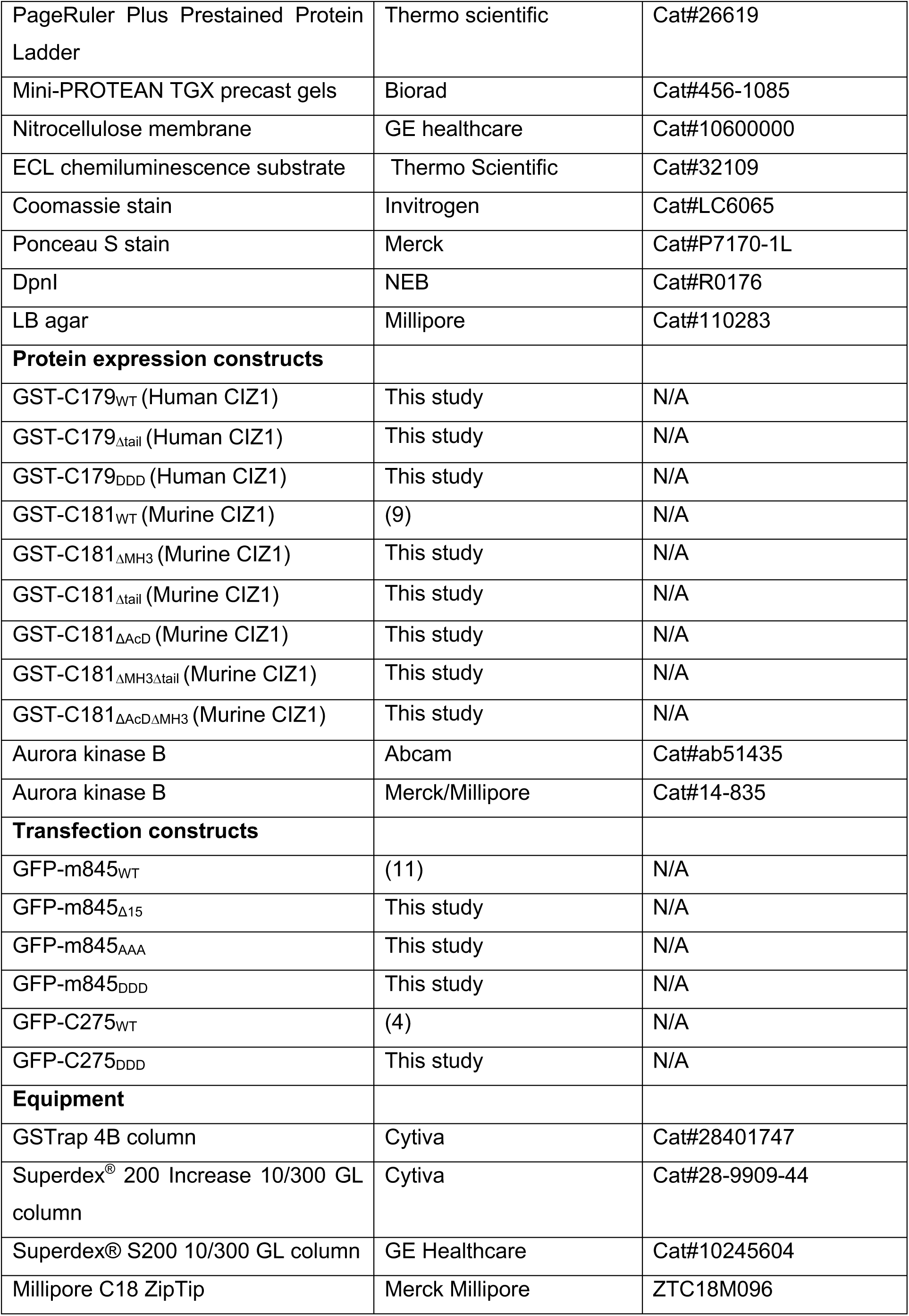
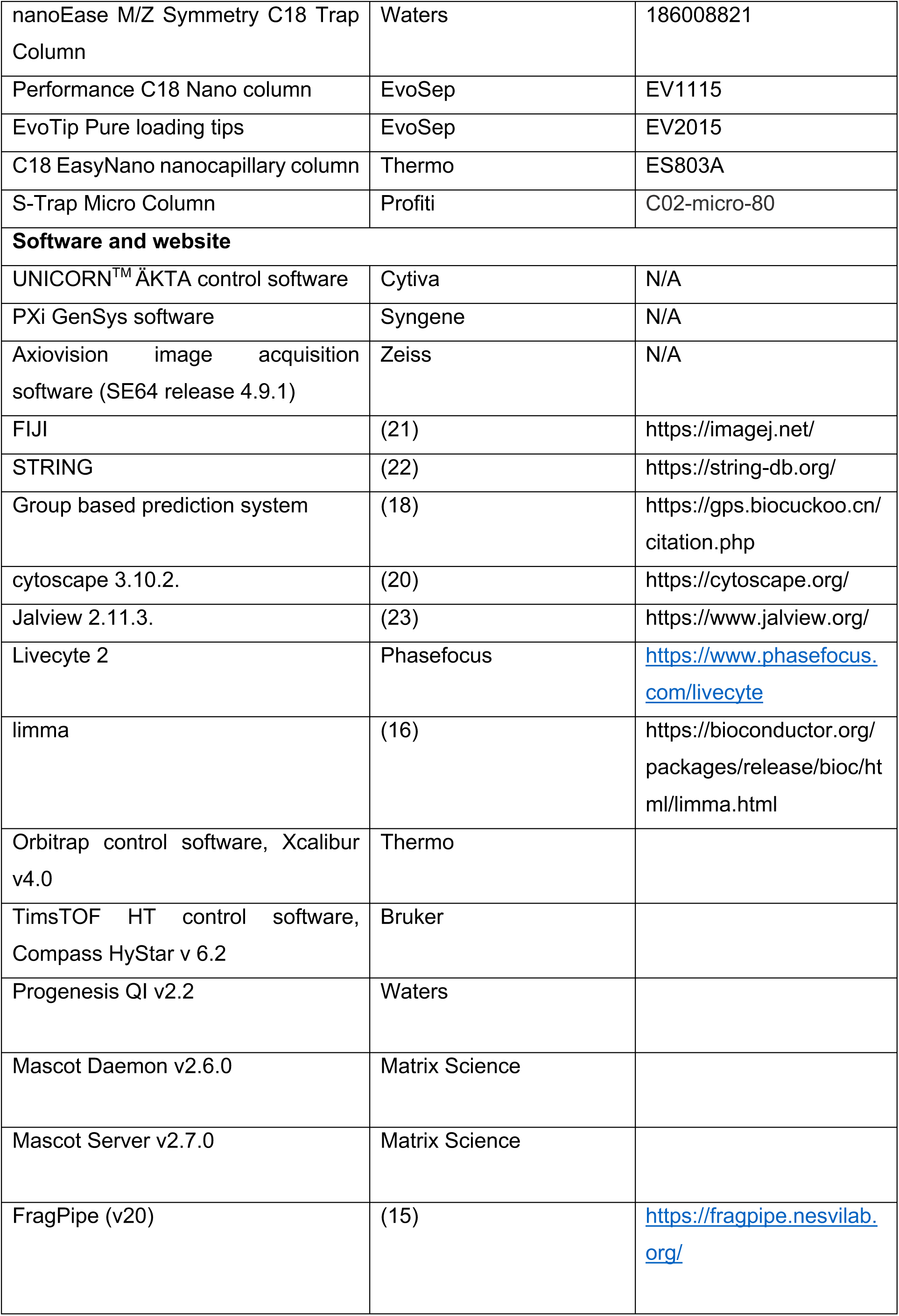
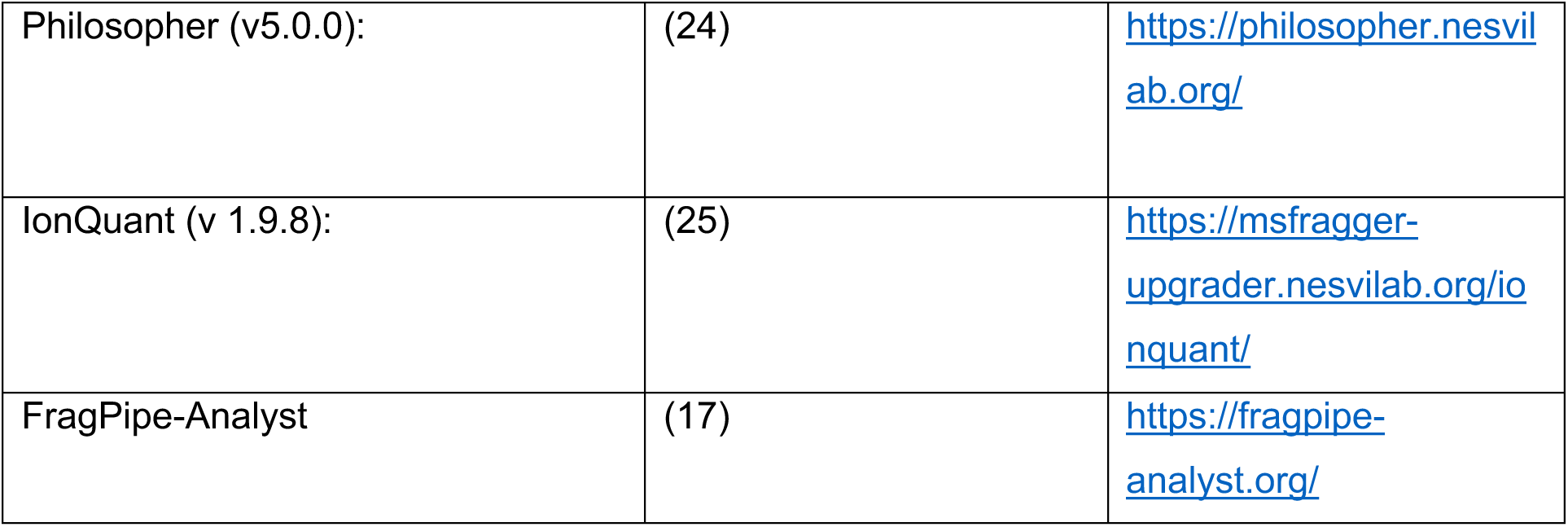
Resources.

### Identification of interaction partners

Protein identification and relative quantification was carried out by label-free mass spectrometry-based proteomic analysis, providing Log2 protein abundance differences (CIZ1/GST control). Differential abundance testing was used to calculate statistical significance between paired sample groups for all identified proteins. Multiple test correction to q-values (adj.p-values) was applied in all cases. Study A compared GST-C179_WT_ to GST control. Study B compared GST-C179_WT_ to GST-C179ϕλ38 and GST control. Study C compared GST-C179_WT_ to GST-C179_DDD_ and GST control. Proteins were digested on-bead with sequencing grade trypsin protease as described (14).

In study A, peptides were desalted with Millipore C_18_ ZipTip before being re-suspended in aqueous 0.1% trifluoroacetic acid (v/v) then loaded onto an mClass nanoflow UPLC system (Waters) equipped with a nanoEaze M/Z Symmetry 100 Å C_18_, 5 µm trap column (180 µm x 20 mm, Waters) and a PepMap, 2 µm, 100 Å, C_18_ EasyNano nanocapillary column (75 μm x 500 mm, Thermo). The trap wash solvent was aqueous 0.05% (v:v) trifluoroacetic acid and the trapping flow rate was 15 µL/min. The trap was washed for 5 min before switching flow to the capillary column. Separation used gradient elution of two solvents: solvent A, aqueous 0.1% (v:v) formic acid; solvent B, acetonitrile containing 0.1% (v:v) formic acid. The flow rate for the capillary column was 300 nL/min and the column temperature was 40°C. The linear multi-step gradient profile was: 3-10% B over 7 mins, 10-35% B over 30 mins, 35-99% B over 5 mins and then proceeded to wash with 99% solvent B for 4 min. The column was returned to initial conditions and re-equilibrated for 15 min before subsequent injections.

The nanoLC system was interfaced with an Orbitrap Fusion Tribrid mass spectrometer (Thermo) with an EasyNano ionisation source (Thermo). Positive ESI-MS and MS^2^ spectra were acquired using Xcalibur software (version 4.0, Thermo). Instrument source settings were: ion spray voltage, 1,900 V; sweep gas, 0 Arb; ion transfer tube temperature; 275°C. MS^1^ spectra were acquired in the Orbitrap with: 120,000 resolution, scan range: *m/z* 375-1,500; AGC target, 4e^5^; max fill time, 100 ms. Data dependant acquisition was performed in top speed mode using a 1 s cycle, selecting the most intense precursors with charge states >1. Easy-IC was used for internal calibration. Dynamic exclusion was performed for 50 s post precursor selection and a minimum threshold for fragmentation was set at 5e^3^. MS^2^ spectra were acquired in the linear ion trap with: scan rate, turbo; quadrupole isolation, 1.6 *m/z*; activation type, HCD; activation energy: 32%; AGC target, 5e^3^; first mass, 110 *m/z*; max fill time, 100 ms. Acquisitions were arranged by Xcalibur to inject ions for all available parallelizable time.

Peak lists in .raw format were imported into Progenesis QI (Version 2.2., Waters) and LC-MS runs aligned. Precursor ion intensities were normalised against total intensity for each acquisition. A combined peak list was exported in .mgf format for database searching against the human subset of the SwissProt database (20,242 sequences; 11,289,677 residues), appended with common proteomic contaminants (116 sequences; 38,371 residues). Mascot Daemon (version 2.6.0, Matrix Science) was used to submit the search to a locally-running copy of the Mascot program (Matrix Science Ltd., version 2.7.0). Search criteria specified: Enzyme, trypsin; Max missed cleavages, 1; Fixed modifications, Carbamidomethyl (C); Variable modifications, Oxidation (M); Peptide tolerance, 3 ppm; MS/MS tolerance, 0.5 Da; Instrument, ESI-TRAP. Peptide identifications were passed through the percolator algorithm to achieve a 1% false discovery rate assessed against a reverse database and individual matches filtered to require minimum expect score of 0.05. The Mascot .XML result file was imported into Progenesis QI and peptide identifications associated with precursor peak areas and matched between runs. Relative protein abundance was calculated using precursor ion areas from non-conflicting unique peptides. Accepted protein quantifications were set to require a minimum of two unique peptide sequences. Statistical testing was performed in Progenesis QI from ArcSinh normalised peptide abundances and ANOVA-derived p-values were converted to multiple test-corrected q-values within Progenesis software.

For studies B and C, LC-MS acquisition was performed using a Bruker TimsTOF HT mass spectrometer. Peptides were loaded onto EvoTip Pure tips for nanoUPLC using an EvoSep One system. A pre-set 60 SPD gradient was used with an 8 cm EvoSep C_18_ Performance column (8 cm x 150 μm x 1.5 μm).

The nanoUPLC system was interfaced to a timsTOF HT mass spectrometer (Bruker) with a CaptiveSpray ionisation source (Source). Positive PASEF-DDA, ESI-MS and MS^2^ spectra were acquired using Compass HyStar software (version 6.2, Bruker). Instrument source settings were: capillary voltage, 1,600 V; dry gas, 3 l/min; dry temperature; 180°C. Spectra were acquired between *m/z* 100-1,700. TIMS settings were: 1/K0 0.6-1.60 V.s/cm^2^; Ramp time, 100 ms; Ramp rate 9.42 Hz. Data dependant acquisition was performed with 10 PASEF ramps and a total cycle time of 1.17 s. An intensity threshold of 2,500 and a target intensity of 20,000 were set with active exclusion applied for 0.4 min post precursor selection. Collision energy was interpolated between 20 eV at 0.6 V.s/cm^2^ to 59 eV at 1.6 V.s/cm^2^.

Data in Bruker .d format were searched using FragPipe (v20.0) (15) against the human subset of UniProt appended with common proteomic contaminants. Search criteria were set as for study A with the exception that, mass tolerance was set to 15 ppm for both MS^1^ and MS^2^. Peptide identifications were processed using philosopher (v5.0.0) to achieve a 1% false discovery rate as assessed against a reverse database. Relative protein quantification was extracted from precursor ion areas using IonQuant (v 1.9.8) with match between runs. Data were filtered to require a minimum of two unique peptides before applying sample minimum imputation. Pairwise statistical comparison was performed using limma (16) via FragPipe-Analyst (17). P-values from pair-wise testing were multiple test corrected to q-value FDR using local and the Hochberg and Benjamini approach. If a protein was deemed to interact with CIZ1, its percentage contribution to total ion area had to be ≥ 2-fold higher than its value when retrieved by GST controls with q ≤ 0.05 considered significant.

### In-vitro phosphorylation

Reactions analysed by western blot were set up using 62.5 ng of recombinant C179_WT_ as substrate with a titration of 0-400 ng Aurora Kinase B (Abcam, ab51435), in 20 mM HEPES pH 7.5, 5 mM MgCl2, 0.1 mM ATP, 1 mM ATP (20 microlitres final volume), and incubated at 30 °C for 30 mins. Reactions were stopped by addition of SDS-PAGE loading buffer. Reactions that were analysed by mass spectrometry contained 5.43 μg recombinant C179_WT_ and 3.15 µg Aurora Kinase B (Merck/Millipore, 14-835), or no kinase control, in 8 mM MOPS/NaOH pH 7.0, 0.2 mM EDTA, 1 % glycerol, 0.02 % v/v beta-mercaptoethanol, 10 mM MgCl2, 0.1 mM ATP, and were incubated at 30°C for 30 min. 5 µg of kinase treated and untreated C179_WT_ were analysed to reveal *in vitro* phosphorylated sites. Treated and non-treated control reactions were split and digested in parallel with either trypsin or Glu-C sequencing grade proteases using an S-Trap (Micro Column, Profiti) mediated proceedure. A 90% aliquot of digested material was enriched for phosphopeptides using MagReSyn titanium dioxide beads. S-Trap digestions and phosphopeptide enrichments were performed as detailed in the manufacturers’ standard protocols. LC-MS data were acquired for both enriched and non-enriched peptides as for *Identification of interaction partners Study B and C*, with the exception that a 100 SPD EvoSep gradient was used. Spectra were searched using FragPipe as detailed above with Glu-C specificity included as applicable and variable phosphorylation of S, T and Y residues also considered. Searches were run with a 1% FDR cut-off. Phosphorylation site localisation confidence was extracted using philosopher (v5.0.0). CIZ1 phosphopeptide identifications were combined for all searches.

### Bioinformatics

Putative kinase sites were mapped on human and murine CIZ1 using data from Group-based Prediction Software GPS 5.0 (18), with additional consensus for AURKB sites from (19).

Analysis of CIZ1 interaction partners was carried out using the STRING database. Networks applied MCL clustering and an inflation parameter of 3, displaying evidence from experimentally determined sources, curated databases and co-expression studies. Outputs were visualised using cytoscape 3.10.2 (20).

### Cloning

Site directed mutagenesis was carried out by PCR, using mutagenic primers listed in Table 2. 4-5 ng of template plasmid was combined with 10-20 ng of primers and CloneAmp HiFi PCR premix (Takara) in 10 µL reactions. Typically, reactions were denatured at 98°C for 30 sec, followed by 18 cycles of 98°C for 10 sec, 55°C for 30 sec, 72°C for 2 min, followed by a final extension of 72°C for 10 min. Products were monitored via agarose gel electrophoresis and successful reactions were chosen for transformation. The PCR products were digested with 2U DpnI (NEB) diluted in 1X CutSmart buffer (NEB) for 1 hr at 37°C. 1 μL of digested PCR products were transformed into 25 μL of DH5α cell suspension (Invitrogen). Cells were thawed and left on ice for 30 min after the addition of DNA, followed by heat shock at 42°C for 45 s with a recovery on ice for 2 min. After an addition of 225 μL SOC broth the reactions were incubated in a shaking incubator for 1 hr and selected by growth on LB agar (Millipore) at 37°C overnight with antibiotic selection. Single colonies were chosen to inoculate 5 mL LB broth for overnight growth in orbital shakers at 37°C 200 rpm and used for plasmid purification via of QIAprep Spin Miniprep Kit (QIAGEN). Sequences were verification by Eurofins Tubeseq service. Plasmids used for protein expression in E. coli were re transformed into BL21 RP cells (Agilent). GFP-m845_Δ15_ (also known as GFP-m830) lacking the terminal 15 amino acids was generated by deletion of a 260bp BamH1 fragment from the C-terminal end of GFP-m845 (11) followed by relegation.

## Results

### Phosphorylation-regulated dissolution of CIZ1-*Xist* assemblies during the cell cycle

During interphase the CIZ1 assemblies that aggregate around Xi chromatin are readily visualised by immunofluorescence microscopy in murine fibroblasts, and other female mouse and human cell types (5-7,10,26,27). However, they become undetectable during the metaphase to anaphase transition, and detectable again approximately 4 hours into G1 phase (Fig.1B,C) (10). This mirrors the cyclical behaviour of *Xist* (28), which is reported to be regulated by the serine/threonine mitotic protein kinase Aurora B (AURKB) (29). Human and murine CIZ1 each encode six canonical AURKB phosphorylation sites of which five are conserved (SFig.1A) - one in the N-terminal half of CIZ1, another in the second Zinc finger, and a cluster of three at the extreme C-terminus (Fig.1D). The C-terminal 38 amino-acids of CIZ1 are predicted to be unstructured in both human and mouse, which possess the same conserved domains (SFig.1B), encoded by the same exons in the same order. This unstructured tail domain and its conserved AURKB site cluster are functionally unexplored.

Initial experiments to explore the cyclical behaviour of CIZ1-Xi assemblies quantified the effect of the AURKB inhibitor barasertib (30) during passage through mitosis. In both D3T3 cells and female primary embryonic fibroblast populations (PEFs at passage 3), the frequency of cells with CIZ1 assemblies remained stable in interphase and prophase (93-94%), dropped to approximately 60% in metaphase, and was almost absent by anaphase (Fig.1E). However, in the presence of barasertib we observed a dose-dependent retention of CIZ1-Xi assemblies in both types of cell in anaphase, increasing from 3% in untreated cells to 84% in the presence of 1µM barasertib. Consistent with the conclusion that phosphorylation promotes dissolution of CIZ1-Xi assemblies, both the broad-spectrum protein phosphatase inhibitor okadaic acid (31) and the concentration-dependent protein phosphatase 1/2A inhibitor tautomycin (32) shifted the regulatory axis and caused dispersal of CIZ1-Xi assemblies in interphase cells within hours (Fig.1F). Dispersal was observed when detected via an epitope in the N-terminal half of CIZ1, and a more complete loss when detected via an epitope in the C-terminal tail of CIZ1 (Fig.1F). Previous work indicates that CIZ1 and *Xist* co-exist within ribonuclear protein particles, and that their accumulation around Xi is co-dependent in fibroblasts (5,6). Consistent with this, *Xist* was also dispersed from interphase cells upon phosphatase inhibition (SFig.1D), as reported previously (29). Together the data argue that release of CIZ1 and *Xist* from Xi is co-regulated by cycles of phosphorylation that involve AURKB. The data also suggest that during interphase either the association between CIZ1 and *Xist*, or the aggregation of CIZ1-*Xist* containing RNPs is actively protected by the action of a phosphatase.

### Direct or indirect effect?

AURKB may phosphorylate CIZ1 to modulate its relationship with *Xist*, however other *Xist*-interaction partners are similarly regulated, and so effects could be indirect. Notably, the DNA-binding domain of HNRNPU/SAFA is phosphorylated by AURKB, and is associated with release of chromosome-associated RNAs during chromosome segregation (33). SAFA normally tethers chromosomal RNAs to chromatin throughout the nucleus in interphase (34), via direct interaction with AT-rich S/MARs (35), and has long been implicated in retention of *Xist* within Xi chromatin territories (36). In our experiments SAFA is lost from the nucleus earlier in mitosis than CIZ1-Xi assemblies, and is also (partially) protected by barasertib in prophase (Fig. 1E). Thus, while AURKB could act directly on CIZ1 to drive its disassembly, these data do not rule out an indirect effect in which phosphorylation of SAFA leads to dispersal of CIZ1 and *Xist*.

### Disruption of the C-terminal tail perturbs nucleus-wide anchoring of CIZ1

To test whether the cluster of three putative AURKB sites in the extreme C-terminus of CIZ1 are functionally relevant, we generated a murine GFP-CIZ1 fusion bearing phosphomimetic mutations in all three sites (GFP-m845_DDD_). When transfected into mammalian cells GFP-m845_WT_ assumes a similar sub-nuclear localisation pattern to the endogenous protein accumulating at both the Xi (in females) and at smaller foci dispersed across the nucleus (in both sexes) (Fig.2A). This requires both N and C terminal domains of CIZ1, is dependent on low complexity prion-like domains that support self-aggregation, and can occur against a CIZ1 null background (9). However, both GFP-m845_DDD,_ and a deletion mutant lacking the terminal 15 amino-acids including the cluster of AURKB sites (GFP-m845_Δ15_), displayed a dramatic phenotype of large sub-nuclear aggregates (Fig.2A,B). In contrast, conversion of the AURKB site cluster to an unphosphorylatable state (GFP-CIZ1_AAA_) did not affect its localisation. Thus, the phosphomimetic mutations drive a dramatic loss of normal function within the C-terminal tail, that affects spatial pattern.

**Fig. 2.**
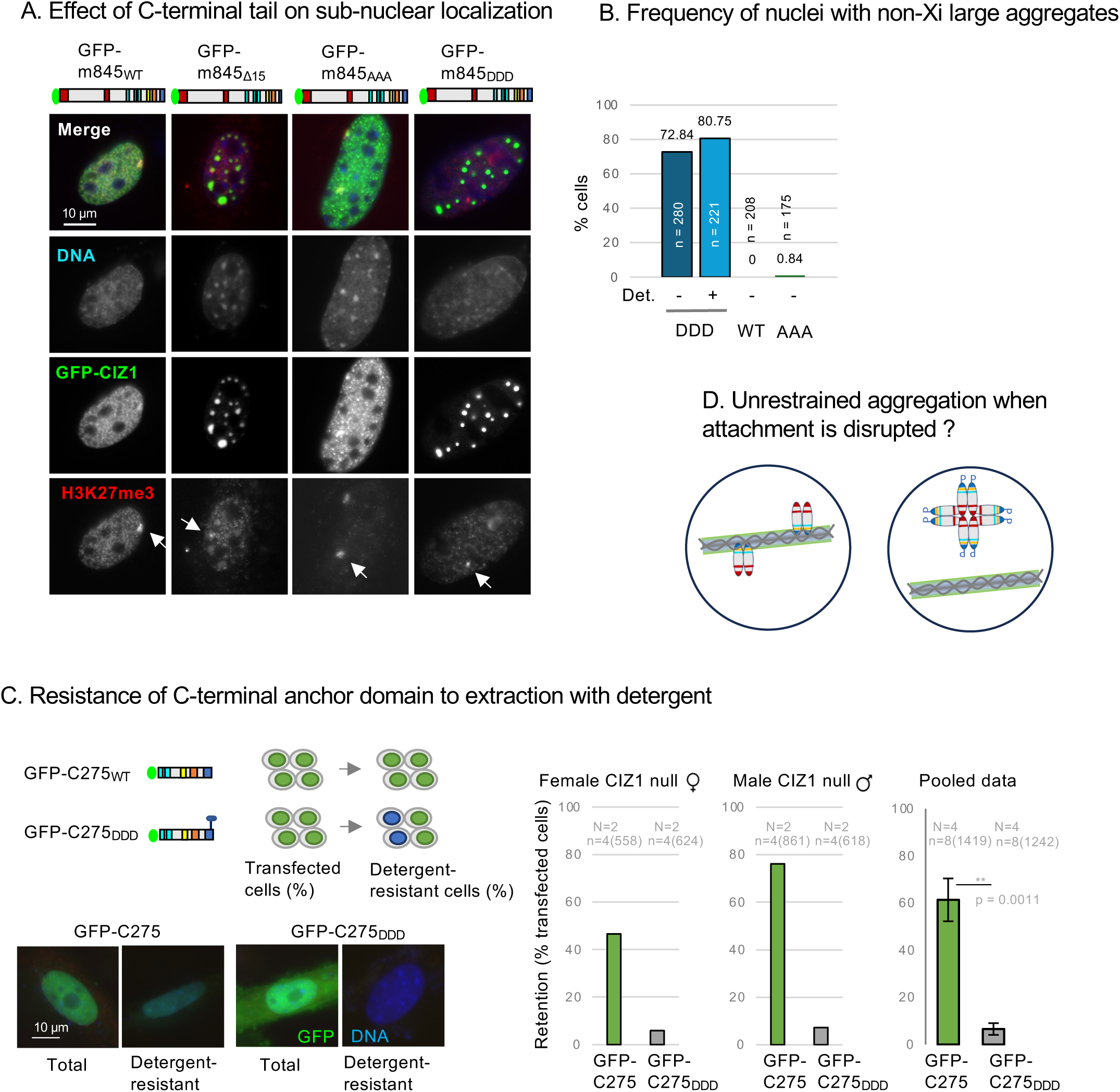
The C-terminal tail specifies nuclear immobilization and limits aggregation. A) Immunofluorescence images showing female murine fibroblasts transfected with full-length mouse GFP-845, or derived constructs m845_Δ15_ (lacking the C-terminal 15 amino acids), m845_DDD_ or m845_AAA_ (green). Cells were co-stained for H3K27me3 (red). DNA is blue, bar is 10 microns. B) Frequency of nuclei containing large (non-Xi) CIZ1 aggregates, shown as percentage of cells transfected with GFP-m845_DDD_ (with and without pre-fixation detergent wash), GFP-m845_AAA_ and EGFP-m845_WT_ control. n = transfected nuclei counted. C) Schematic showing the C-terminal portion of CIZ1, GFP-mC275 and derived GFP-mC275 _DDD_, and their use in 48 hour transient expression experiments to assess ability to assemble into detergent-resistant structures (4). Below, example images of transfected nuclei. Histograms show retention frequency in male (N=2) and female (N=2) CIZ1 null primary embryonic fibroblasts that were transfected (green). n denotes technical replicates for each cell population, with number of nuclei scored in parentheses. D) Illustration displaying the effect of AURKB site cluster phosphomimic (P) on CIZ1’s association with chromatin, and associated detergent-resistant nuclear structures.

When tested in the context of the C-terminal nuclear matrix anchor domain alone (GFP-C275) which lacks the PLDs and other N-terminal sequences, GFP-C275_DDD_ revealed failure to become immobilized by attachment to insoluble nuclear structures. Like GFP-C275 (4,9), GFP-C275_DDD_ was expressed and imported into the nucleus (where it did not form large aggregates), but its resistance to extraction by detergent was reduced from 61.3% resistant nuclei to 6.6%. This was tested in CIZ1 null cells to avoid complications arising from possible association with the endogenous protein, and in both male and female cells to enable wider interpretation of the results beyond possible interaction with female-specific *Xist* (Fig.2C). Together these observations suggest the C-terminal tail supports anchorage of CIZ1 in the nucleus, and are also consistent with the idea that this limits the PLD-driven tendency of CIZ1 to self-aggregate into large assemblies (Fig.2D).

### Modification of C-terminal AURKB sites in metaphase

Anti-CIZ1 peptide antibody, raised against sequences in the C-terminal tail in which the AURKB sites reside (Fig.1D, SFig1C, ‘tail antibody’) does not detect ectopic full length GFP-m845_DDD_ in cell lysates by western blot, while an N-terminal anti-CIZ1 antibody does (Fig.3A). This shows that its epitope is affected by the mutations. When applied to whole cell lysates from synchronised cells expressing only endogenous CIZ1, it also strikingly fails to robustly detect the C-terminal epitope in some contexts. Cell populations arrested in mitosis, or cycling populations treated with the phosphatase inhibitor okadaic acid, were poorly detected compared to N-terminal epitopes (Fig. 3B), and compared to untreated cycling populations or those arrested in S phase, in which N and C terminal epitopes are both available. This suggests that CIZ1 is modified by phosphorylation of C-terminal tail sequences in mitosis (Fig.3C), and that the event can be reported via this phospho-sensitive anti-tail antibody.

**Fig. 3.**
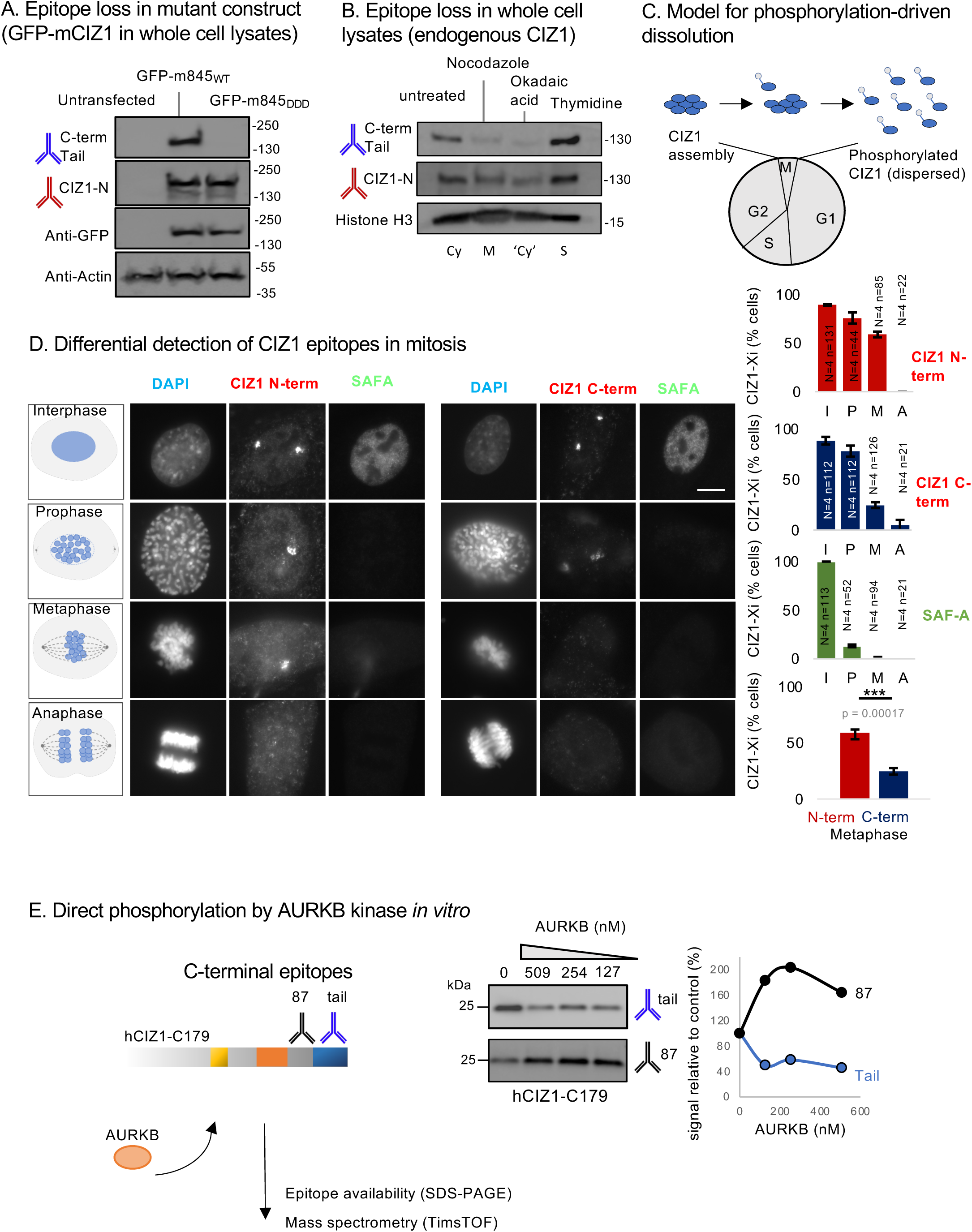
AURKB site modification in mitosis. A) Western blot showing denatured proteins in whole cell lysates collected from untransfected D3T3 cells, or populations expressing full length mouse GFP-m845_WT_ or GFP-m845_DDD_, after immunostaining for CIZ1-N or CIZ1-C (tail epitope), or b-actin, and GFP as indicated. B) Western blot showing denatured endogenous proteins in chemically treated D3T3 cells, to achieve cell cycle enrichment in mitosis (M, nocodazole), S phase (S, thymidine), or a phosphatase-suppressed state (okadaic acid). Immunoblotting for C-terminal tail and N-terminal CIZ1 indicates a reduction in tail epitope, compared to untreated cells, during arrest in metaphase, or after phosphatase inhibition. Histone H3 is shown as a loading control. Cy, cycling. C) Illustration showing data interpretation in which the CIZ1 tail AURKB site cluster is phosphorylated in mitosis, driving dispersal of CIZ1 from Xi assemblies. D) Female D3T3 cells in stages of mitosis as indicated, immuno-stained for CIZ1-N or CIZ1-C and co-stained for SAFA. Right, histograms show frequency of retention in interphase (I), prophase (P), metaphase (M) or anaphase (A), where N indicates replicate analyses and n nuclei scored. Lower, by metaphase CIZ1-N and CIZ1-C are significantly different (p<0.00017), students t-test. E) Experimental overview of *in vitro* kinase reactions using purified recombinant human CIZ1 C-terminal fragment C179 and purified AURKB. Middle, products analysed by western blot with C-terminal CIZ1 epitope-defined antibodies, showing changes in reactivity in response to exposure to increasing concentrations of AURKB kinase. Graph shows band intensities relative to untreated C179 control. Products were also analysed by mass-spectrometry (SFig.2C).

N-terminal and C-terminal antibodies also report different dynamics during mitosis when used for immunostaining of endogenous CIZ1 (Fig.3D). Though the frequency of cells with CIZ1-marked Xi’s is similar in interphase populations (88.5% C-term, 89.1% N-term) and prophase populations (78.1% C-term, 75.8% N-term), in metaphase nuclei the C-terminal epitope is significantly less frequent (24.6% C-term, 58.8% cells N-term). This was confirmed using a different (goat) anti-tail antibody in co-staining experiments (SFig.2A), and suggests that the tail epitope is modified before assembly dissolution. Together the data support the idea that CIZ1 is normally phosphorylated in its C-terminal AURKB sites between prophase and metaphase, likely contributing to dissolution of CIZ1-Xi assemblies.

### AURKB directly phosphorylates the CIZ1 C-terminus

To confirm that CIZ1 can be directly phosphorylated by AURKB, recombinant human CIZ1-C179 protein was incubated with recombinant AURKB, and the impact assessed using phosphosensitive C-terminal tail antibody by western blot (Fig.3E). Compared to mock reactions, AURKB reduced the reactivity of the C-terminal tail antibody by 54%, supporting direct modification of the tail. Interestingly, AURKB exposure also increased accessibility to the C-terminal mAb 87 epitope by 63%, despite its epitope being located upstream of the putative kinase sites in the linear protein sequence (Fig.1D). The products of AURKB exposure were further probed in a phosphoproteomic analysis, which identified ten high confidence targets in recombinant hC179 after exposure to AURKB, six within the C-terminal tail (SFig2B,C). Of the three clustered conserved AURKB sites that were mutated only the terminal residue (Thr 898) was among the high confidence sites. Nevertheless, these data confirm that CIZ1 is a potential substrate of AURKB.

### C-terminal interaction partners are enriched in chromatin and nuclear matrix proteins

To explore the interactions that normally anchor CIZ1 within the nucleus, and which may be interrupted by phosphorylation of the AURKB site cluster in the CIZ1 tail, we first established a protein interaction network. Three independent studies, using the C-terminal 179 amino acids of human CIZ1 with an N-terminal GST tag, retrieved high-confidence interaction partners from nuclear extracts prepared from cycling HeLa cells (Supplemental Dataset 1). Across the three studies, 118, 263 and 187 proteins were identified that prefer GST-hC179 compared to GST alone (Fig.4A), of which 56 were identified in all three studies, and a further 78 in two of the three studies (Fig.4B, Supplemental Dataset 2).

**Fig. 4.**
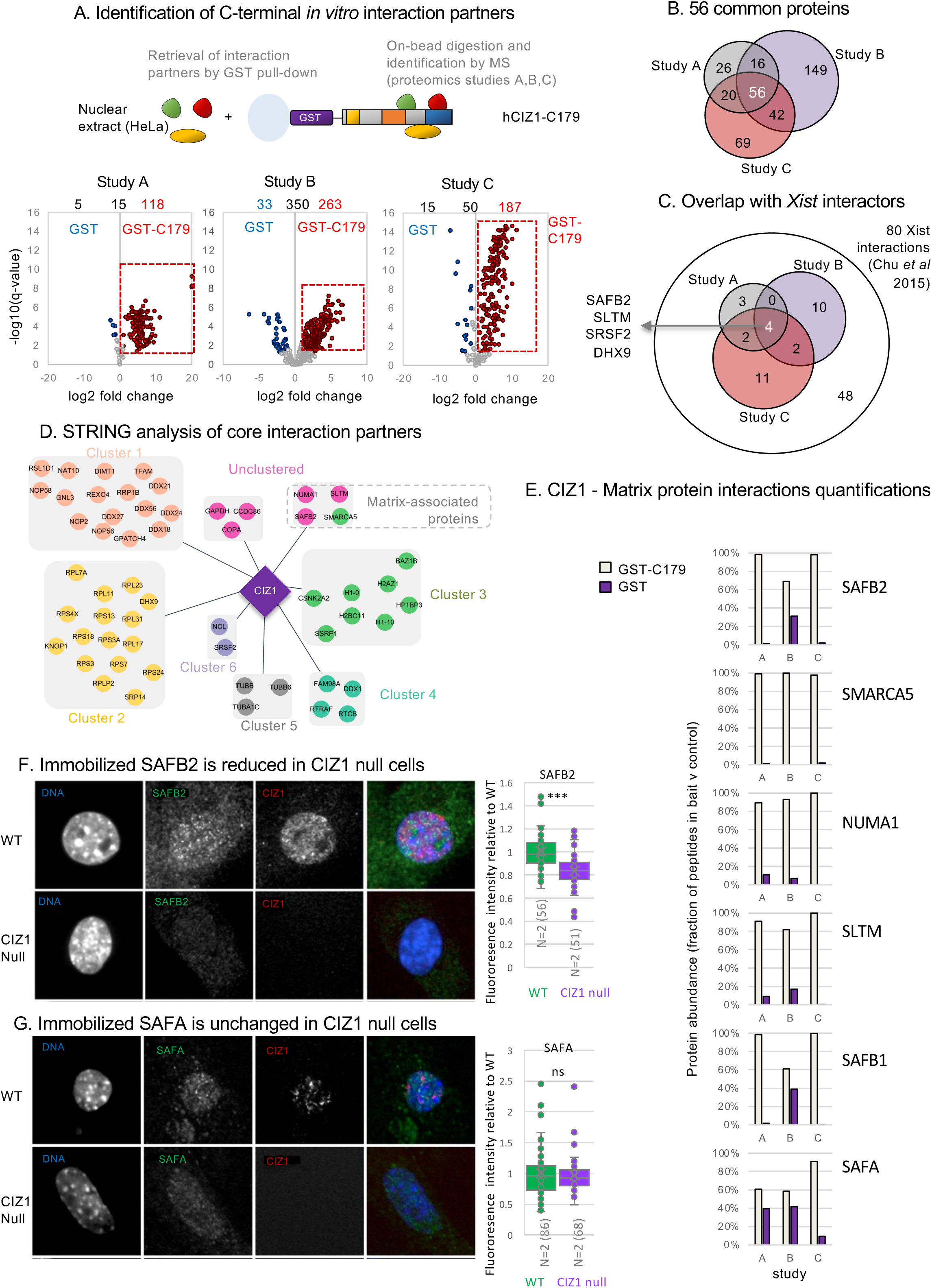
CIZ1 C-terminal interaction partners. A) Overview of nuclear protein interaction studies using mammalian cell nuclear extracts from Hela cells and recombinant GST-tagged hCIZ1 C-terminal fragment C179. Domains coloured as in Fig.1D. Below, volcano plots showing protein interaction partners identified with high confidence in three independent studies, and their relative retrieval by GST-hC179 compared to GST control. Significance (-log10 FDR q-value) is plotted against log2 fold change (FC), derived from n=4 replicates in each case. Significant interaction partners are ≥ 2 fold more abundant in CIZ1-retrived samples compared to GST samples, at q-value ≤ 0.05. Data is given in Supplemental Dataset 1. B) Venn diagram showing common interaction partners between three independent studies. The 56 core interaction partners are listed in Supplemental Dataset 2. C) CIZ1 interaction partners (red segment) in common with 80 *Xist* interactors identified by CHIRP-MS (37). Venn diagram indicates those that interact with CIZ1 in all three of our studies. D) Simplified STRING diagram showing 56 core CIZ1 interaction partners, clustered using MCL clustering. Three unclustered proteins (dark pink), and one chromatin protein (green) are nuclear matrix -associated proteins (Supplemental Dataset 2). E) Individual abundance (mean of four replicates) across three studies for four nuclear matrix proteins in the core 56 interaction list (SAFB2, SMARCA5, NUMA1, SLTM) and two that appear in at least one of the studies (SAFA, SAFB1). Histograms show fraction of high confidence peptide in bait and control for each study. F) Example immunofluorescence images of SAFB2 (green) in cycling WT and CIZ1 null PEFs, co-stained for CIZ1-N (red), and Dapi (blue). Box and whisker plots show intensity measures derived from two independent primary cell populations (N) for each genotype. n=number of nuclei measured. Comparison is by t-test where *** denotes p<0.001, and indicates a significant reduction of bound SAFB2 epitope in CIZ1 null cells. G) As in F but for SAFA, which is not significantly changed in CIZ1 null cells.

*Xist* lncRNA has been extensively probed to identify protein interaction partners in different cells types, using a range of methods (37-40). We compared our core list of 56 human CIZ1 interaction partners, with the 81 proteins retrieved from four stem cell lines by *Xist* CHIRP-MS (37). Excluding CIZ1 itself, four proteins were common (DHX9, SAFB2, SLTM, SRSF2) across all experiments, but we observed a further 28 across at least one of our three studies (Fig.4C, Supplemental Dataset 2), including the nuclear matrix proteins SAFA, Matrin-3 and HNRNPK.

STRING network analysis (22) of the 56 common proteins separated 50 into six clusters (Fig.4D), and Gene Set Enrichment Analysis (GSEA) within the STRING platform assigned functional groups (Supplemental Dataset 2). Clusters 1 and 2 contain proteins linked with RNA binding and RNA processing, and include DHX9, which was previously reported to interact with the N-terminal part of CIZ1 (41), plus five other DEAD-Box helicases. All CIZ1 interaction partners in cluster 3 belong to the cellular component chromatin (GO:0000785), including two nucleosomal histone variants and two variants of linker histone H1, as well as several nucleosome remodelling factors. The six unclustered proteins returned no GO processes or function, however they include three nuclear matrix associated proteins - NUMA1 (42), and the Scaffold attachment factor B family members SAFB2 and SLTM (SAFB-like transcription modulator) (43,44). SAFB2 was previously identified in the bioplex network as a potential CIZ1 interaction partner (45). SMARCA5, a component of the ‘Imitation SWItch’ (ISWI) chromatin remodelling complex, assigned to cluster 3 is also a nuclear matrix-associated protein (Fig.4D,E).

To probe possible functional relationships between CIZ1 and this class of proteins we looked at their sub-nuclear location and retention in CIZ1 null PEFs. Despite no significant difference in transcript levels compared to WT PEFs (SFig.3A), significant differences in their nuclear retention were observed for some. In interphase cells, SAFB2 is reduced, SAFA is unchanged, and SMARCA5 is increased (Fig.4F,G, SFig.3B). Thus, SAFB2 may, in part, be dependent on CIZ1 for accurate assembly.

### CIZ1 null cells display segregation defects

In the same primary cell types we observed CIZ1-related mitotic deficiencies. Two independent CIZ1 null PEF populations showed increased frequency of spindle abnormalities compared to WT (SFig.3C), and a significant increase in the frequency of aborted mitoses, as revealed by live cell tracking of parallel populations (SFig.3D,E,F). While the requirement for CIZ1 for high-fidelity spindle function is not yet understood this does implicate it, and possibly its binding partners, in the accurate execution of mitosis.

### Phosphomimetic CIZ1 mutants have impaired interaction with RNA binding proteins

The nuclear matrix proteins with which CIZ1 interacts are candidate factors that could support its localisation and anchorage within the nucleus, and which may be sensitive to AURKB phosphorylation during CIZ1-Xi assembly dissolution in mitosis. We tested this by evaluating the effect of deleting the C-terminal 38 amino acids (C179_Δtail_ in study B), or of converting the three clustered AURKB sites to aspartic acid (C179_DDD_ in study C), which revealed unexpected results (Fig.5A,B,C detailed in Supplemental dataset 2). While C179_DDD_ lost 37 proteins (including 10 of the core 56 interaction partners), complete deletion of the tail region resulted in no loss of any core proteins that met the significance threshold (FDR q<0.05, log2 fold change >1). In fact, many are increased, including five core interaction partners (BAZ1B, CCDC86, DDX24, NUMA1 and RPL11), which contribute to a total of 94 proteins that prefer C179_Δtail_ compared to WT. One potential explanation for the emergence of new interaction partners is that the 38 amino acids that make up the unstructured tail region could block interaction sites located elsewhere.

**Fig. 5.**
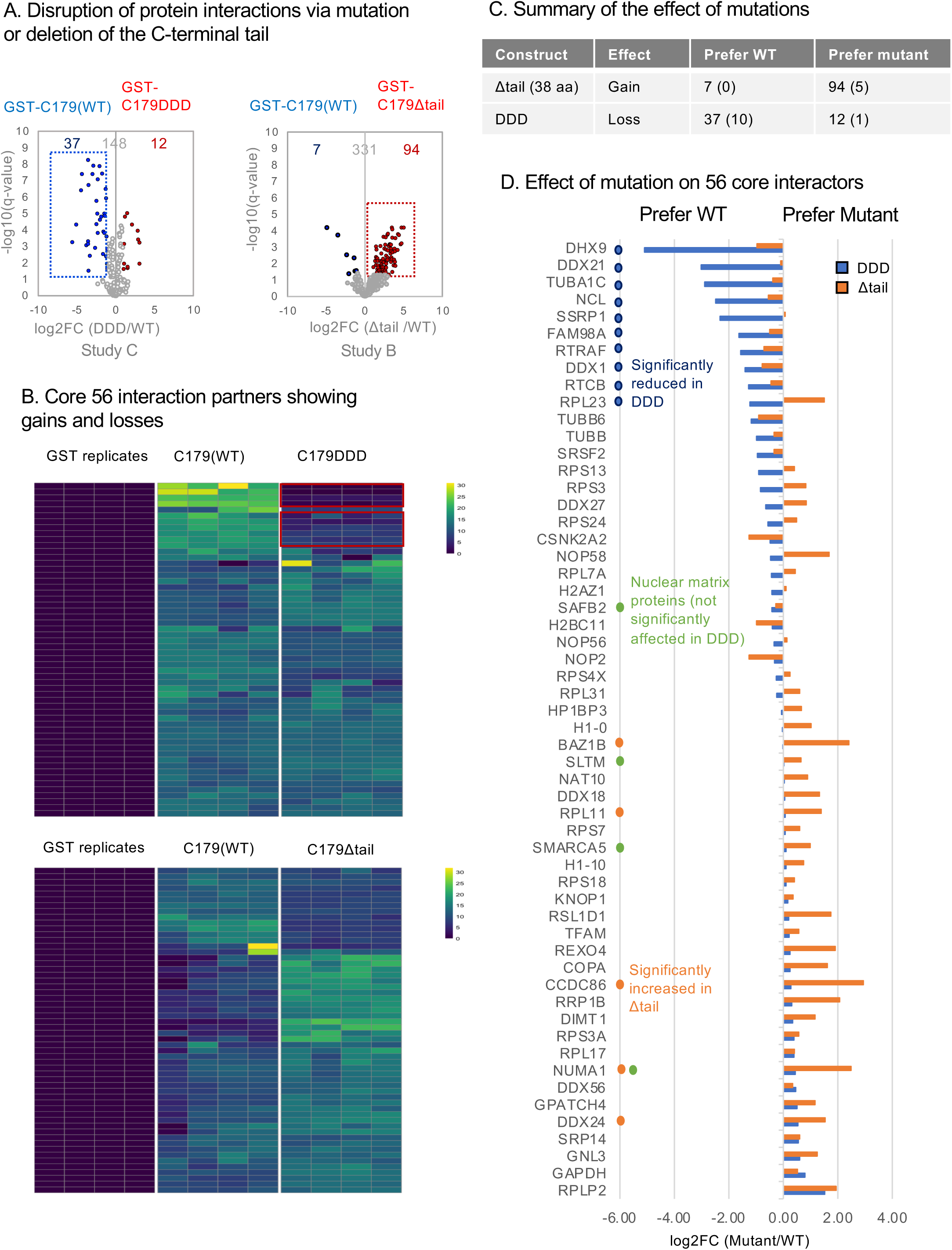
Effect of mutation on CIZ1 C-terminal interaction partners. A) Volcano plots displaying protein interaction partners, comparing WT human C179 with C179_DDD_ (left) or C179_Δtail_ (right), in independent studies. The majority of proteins were unaffected. Those significantly increased or decreased are highlighted (log2 fold change +/- 1, q ≤ 0.05). See also Supplemental Dataset 2. B) Heatmaps showing peptide abundance of the core 56 interaction partners retrieved in control (GST), WT (C179) and mutant reactions, as indicated (n=4 replicates). Upper, 10 proteins which are significantly reduced in C179_DDD_ compared to WT are indicated. Lower, of the core 56 only 5 are significantly changed (see D). C) Summary table describing effects of mutations on study specific interaction partners, and core 56 interaction partners (in parentheses). D) Plot displays log2FC of core 56 proteins for C179_DDD_ (blue) and C179 _Δtail_ (orange), with identities. Ten proteins that are significantly reduced upon phosphomimetic mutation are labelled. Nuclear matrix proteins SAFB2, SLTM, SMARCA5 are not affected, however NUMA1 was increased in C179 _Δtail_.

Of the four nuclear matrix proteins in the core 56 interaction partners, none were lost (Fig.5D). Thus, regulated dissolution of CIZ1 assemblies, via phosphorylation of the C-terminal AURKB site cluster, does not appear to be achieved via dissociation from other nuclear matrix proteins.

GSEA of the 37 proteins whose binding is significantly diminished in study C, revealed that 33 are associated with Gene Ontology (GO) molecular function RNA binding (GO:0003723, FDR 1.39E-27), of which 17 have RNA splicing function (GO:0008380, FDR 1.15E-15), while the 10 that are part of the core 56 interaction list include the entirety of cluster 4 tRNA-splicing ligase complex (Fig.5D, Supplemental dataset 2). Thus, phosphomimetic mutation of CIZ1 does affect *in vitro* protein interactions, but 89% of those affected are themselves RNA binding proteins. These could undergo direct protein-protein interaction with CIZ1, or could have been retrieved via indirect interaction mediated by RNA. This lead us to ask whether phosphorylation of the C-terminal AURKB site cluster could cause dissolution of CIZ1-Xi assemblies by driving dissociation from direct interaction with RNA.

### CIZ1 exists as a stable dimer with MH3 domain interface

To allow evaluation of the effect of the C-terminal tail on interaction with RNA, we expressed, purified and partially characterised a series of human (C179) and mouse (C181)-derived proteins (Fig.6A, SFig.4A). Size exclusion chromatography (SEC) indicated that WT forms of both species exist in a stable multimeric state (SFig.4). SEC coupled with multiple angle laser light scattering (SEC-MALLs) confirmed that both are homodimers with molecular weight estimates of approximately 44 kDa, consistent with dimerization of the ∼20 kDa monomer (Fig.6B,C,D). Deletion of the C-terminal 38/37 amino acids from either the mouse or human proteins, or phosphomimetic (DDD) mutation of the human protein had minimal effect on elution time in SEC, and SEC-MALLS confirmed human hC179_Δtail_ still to be a stable dimer (Fig.6B). However, deletion of the MH3 domain significantly delayed SEC elution, and SEC-MALLs verified the ΔMH3 mutant is a monomer (Fig.6C,D), indicating that the MH3 domain is required for dimerization. Modelling of secondary structure of human and murine CIZ1 MH3 domain using AlphaFold2 (46,47) to form a dimeric entity, highlighted a conserved β-strand as the putative dimerization interface (Fig.6E). Individual full SEC and SEC-MALLs traces are given in SFig.4.

**Fig. 6.**
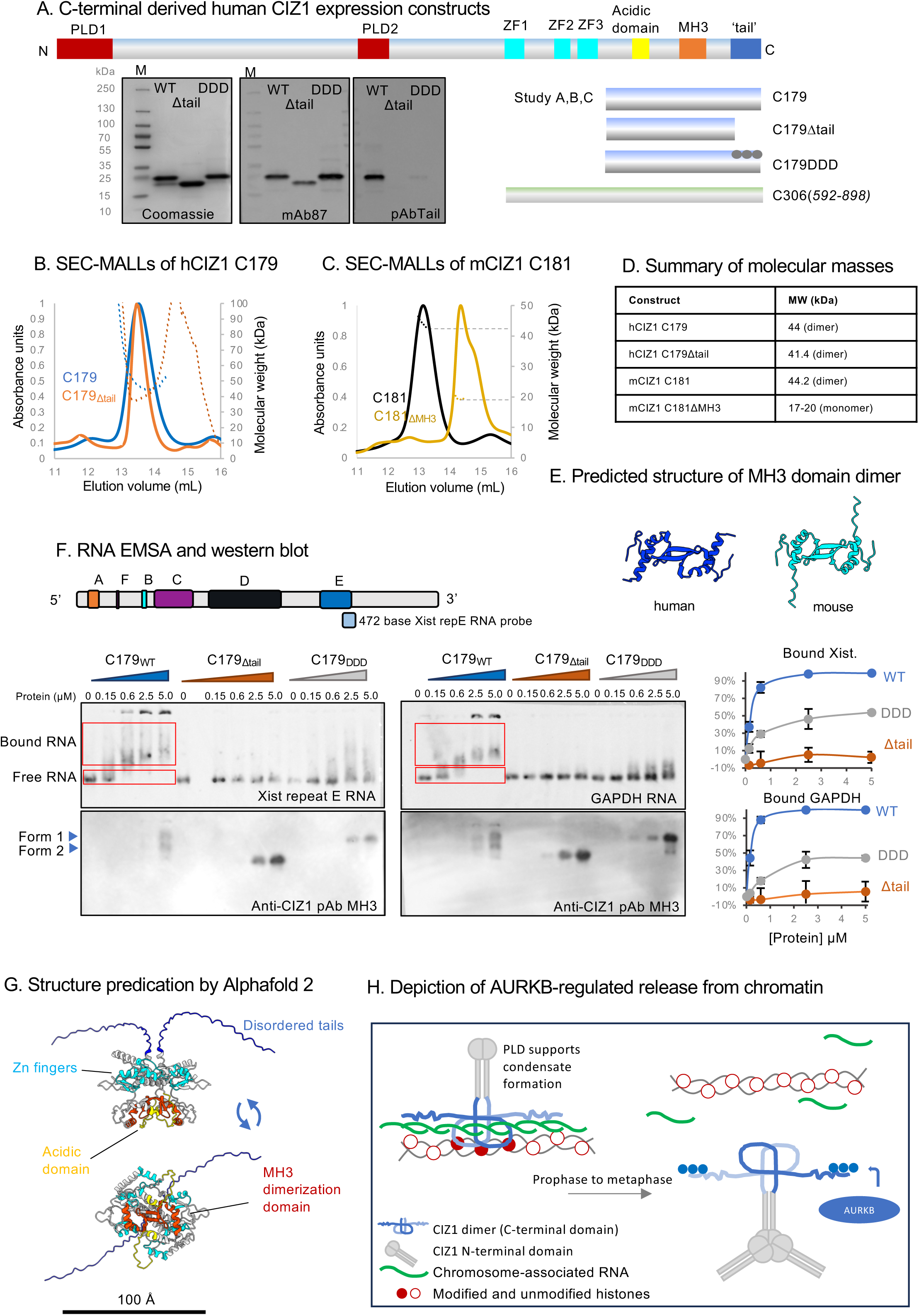
Interaction between CIZ1 dimer and RNA is regulated by AURKB sites in the C-terminal tail. A) Schematic of h/m CIZ1 showing conserved domains, in yellow (acidic domain), orange (MH3 dimerization domain) and blue (unstructured tail h37/m38 C-terminal amino-acids). Below, human C-terminal (C179) fragments, and derived mutants used as bait fragments in interaction studies, including C179_Δtail_ and phosphomimic C179_DDD_. Below, C-terminal fragment encompassing the Zn finger motifs used for modelling (green, C305). Left, SDS-PAGE gels showing purified protein preparations stained with Coomassie blue, or probed with anti-CIZ1 mAb 87 which recognises all three proteins, or anti-CIZ1 tail pAb which recognises an epitope deleted in C179_Δtail_ and mutated in C179_DDD_. B) Size exclusion chromatogram with Multi Angle Laser Light Scattering (SEC-MALLS), showing normalised UV absorbance at 280nm, and molar mass (dotted line), for human CIZ1-C179 (blue) and human CIZ1-C179_Δtail_ (orange). C) SEC-MALLS chromatogram showing normalised UV absorbance at 280nm and molar mass (dotted line) for equivalent murine fragment C181 (black), and derived deletion mutant lacking the matrin 3 homology domain (C181_ΔMH3_, yellow). D) Summary of measured molecular masses, indicating that the C-terminal fragment forms a stable dimer, that is dependent on the MH3 domain but not the tail region. E) AlphaFold dimer structure predictions of MH3 domain, showing human CIZ1 aa779 to 838 uniprot Q9ULV3-1 (blue), and murine CIZ1 aa725 to 785 uniprot Q8VEH2 (cyan). The domain forms a tight dimer with monomer-monomer interactions involving main chain hydrogen bonding between β-strands of the two MH3-type Zn finger motifs. F) Example electrophoretic mobility shift assays (EMSA) showing the effect of C179, C179_Δtail_ and C179_DDD_ on the mobility of digoxygenin (DIG) labelled *Xist* repeat E RNA probe (left, 0.66 nM) or GAPDH RNA (right, 0.65 nM). Below, immunoblots of EMSA membranes using CIZ1 anti-MH3 domain antibody. Above, murine *Xist* structure (63) and the derived *Xist* repeat E RNA probe used in EMSAs. Right, quantification of binding based on the fraction of shifted probe, derived from three replicate experiments (see also SFig4). Graphs show means ± SEM. G) AlphaFold-Multimer (46,47) structure prediction of human C-terminal aa 592-898 (hC306), showing the highest-ranking prediction, in which the acidic domains (yellow) are exposed and the unstructured tails (blue) extend from the core. H) Model, depicting CIZ1 homodimers interacting with chromosome-associated RNAs via its C-terminal tails, with N-terminal PLD domains available for association with other proteins or other RNAs (left). Right, shows AURKB-mediated phosphorylation driving release from chromosome-associated RNA. *In vitro* in interphase this results in PLD-driven CIZ1 aggregation.

### Direct interaction with RNA via the C-terminal tail

Electrophoretic mobility shift assays (EMSA) have shown that murine CIZ1 (C181) can interact directly with RNAs, including *Xist* repeat E and GAPDH (9). Here, we show that this capability is lost in the ΔMH3 mutant (SFig.4C), suggesting that dimerization may be involved in the creation of a stable RNA interaction interface. Furthermore, using the human WT version (C179), we confirmed direct interaction with RNA and showed this to be mediated by the tails (Fig.6F). WT C179 shifts 99% of *Xist* probe when used at 5µM, while the phosphomimetic mutant (C179_DDD_) shifts only 54% and the tail deletion mutant (C179_Δtail_) only 2%, under the same conditions. Similar results were returned when GAPDH RNA was used. Together these results confirm direct interaction with RNA, identify the unstructured C-terminal tail region of CIZ1 as an RNA-binding sequence, and show that interaction is likely modulated by phosphorylation of sites that are subject to cell-cycle dependent regulation.

### Modelling the CIZ1 dimer

In a longer polypeptide (human C306, Fig.6A), encompassing the three conserved zinc fingers, the MH3 domains forms a tight dimer, with exposed acidic patches and unstructured C-terminal ‘tails’ that protrude from the core, with the AURKB sites available for interaction (Fig.6G). Thus, AlphaFold 2 predictions are consistent with our experimental findings, and contribute to a model in which the C-terminal tails of CIZ1 are normally embedded in chromatin, via direct interaction with chromosome-associated RNAs, including *Xist*. Dissociation, driven by AURKB in mitosis, leads to CIZ1 assembly dissolution and scheduled exposure of underlying chromatin (Fig.6H).

## Discussion

Early concepts of a stable network of RNA-dependent proteins that form a nucleus-wide matrix have shifted to a more dynamic model in which RNAs seed localised protein assemblies (2). Non-coding RNAs are integral to the formation of a range of membrane-less structures within the nucleus, including the nucleolus, the Xi territory and other RNA-chromatin compartments (48), and more recently 3D maps have revealed how ncRNA-mediated recruitment of proteins to sites of transcription can regulate chromatin state (49). However, the extent to which these RNA-localised functional assemblies are cross-linked into a wider network is not known, and may in fact vary with cell type and cell state. Most of the analysis described here is carried out in cycling primary fibroblasts, in which the CIZ1-RNA assemblies that gather around the Xi are rapidly dispersed and rebuilt each cell cycle. This study does not necessarily shed light therefore on their characteristics in differentiated and largely quiescent tissues.

Together with earlier analysis of the low-complexity prion-like domains (PLDs) in the N-terminal half of CIZ1 (9), the present study validates CIZ1 as a multivalent RNA-binding protein. We show that CIZ1 molecules form stable homodimers *in vitro* dependent on a highly structured MH3 domain, and that while both N- and C-terminal RNA interaction interfaces are required to form large protein assemblies at the Xi (9), modulation of the one in the dual C-terminal tails is sufficient to disrupt accumulation at the Xi, as well as anchorage elsewhere in the nucleus. Moreover, under conditions in which anchorage is impaired, CIZ1 coalesces into abnormally large aggregates, likely driven by its PLDs. This suggests that anchorage creates fixed points of coalescence, that restricts and limit the number of molecules that are free to aggregate at any one location.

In addition to those that gather within the Xi territory, CIZ1 forms more than a hundred smaller assemblies across the nucleus likely seeded by lncRNAs other than *Xist*. While the full scale of its RNA interactome is not known, altered expression of CIZ1 in early-stage breast cancers has been shown to disproportionally affect the levels of lncRNAs, so defining a candidate human cohort (10). Thus, the phosphorylation-regulated dissociation of CIZ1 from RNA reported in the present study may be relevant to autosomal assemblies in addition to those at the Xi. Certainly, the consequences of CIZ1 loss or disruption are felt across the genome (8,10), and its direct interaction with RNA is not limited to *Xist*. We postulate that AURKB phosphorylation could drive a wider dissociation of CIZ1-RNA contacts across the nucleus, via a direct effect on its tails.

Phosphorylation regulates multiple steps in mitosis from spindle formation (50) to nuclear envelope deconstruction (51-53) and its deregulation is implicated in disease states. AURKB is overexpressed in lung (54), breast (55) and prostate cancers (56), and is associated with chromosome mis-segregation leading to aneuploidy (57). The nuclear matrix protein SAFA is also phosphorylated by AURKB during mitosis, and failure of AURKB-dependent disassembly, and removal of chromatin-associated RNAs during prophase is mechanistically implicated in elevated rates of anaphase segregation defects, and reduced fidelity of chromosome segregation (33). In our studies, while we have shown that CIZ1 is required for high-fidelity chromatid division, it is not yet clear how these observations relate to the regulated behaviour of SAFA. Our data does not directly implicate failure to remove chromatin-associated RNAs, however the links between CIZ1 and cancer both in murine models (5,58), and *in vivo* in humans (10,59-62), should be considered in the light of aberrant mitoses, as well as the defects in epigenetic control of gene expression that we reported previously (8).

## Data Availability

Raw mass spectrometry data and proteomic results files are referenced in ProteomeXchange (PXD067610) and can be obtained from MassIVE (MSV000098911, [doi:10.25345/C51Z4259D]).

## Acknowledgements

We thank Andrew Leach for help with SEC-MALs, Maria Chechik for guidance on protein expression, Emma Stewart and Louisa Williamson for fibroblast isolation, genotyping and additional FISH analysis. We are also grateful to former students whose work did not make it into this paper, including Oliver Smith and Izak Vary. The York Centre of Excellence in Mass Spectrometry was supported by Science City York with funds from the Northern Way Initiative and by Engineering and Physical Sciences Research Council grants EP/K039660/1 and EP/M028127/1. This study was funded by BBSRC doctoral training fellowship BB/T007222/1 to LB, the Georgina Gatenby PhD scholarship to GT, and in part by Royal Society Leverhulme Trust Fellowship SRF\R1\191028 to DC, with additional support from Wellcome Trust grant 224665 to AAA, and Medical Research Council grant MR/V029088/1 to DC.

## Author contributions

LB, GT, DC designed experiments.

LB, GT, MT, NS, BG, MB, CB, KC, WD, EN performed experiments.

LB, GT, AD, JA, DC analysed experiments.

JA, LB, SS, EG established methodologies and created materials.

AA, DC supervised the project.

LB and DC wrote the paper.

## Supplemental information

**SFig.1.**
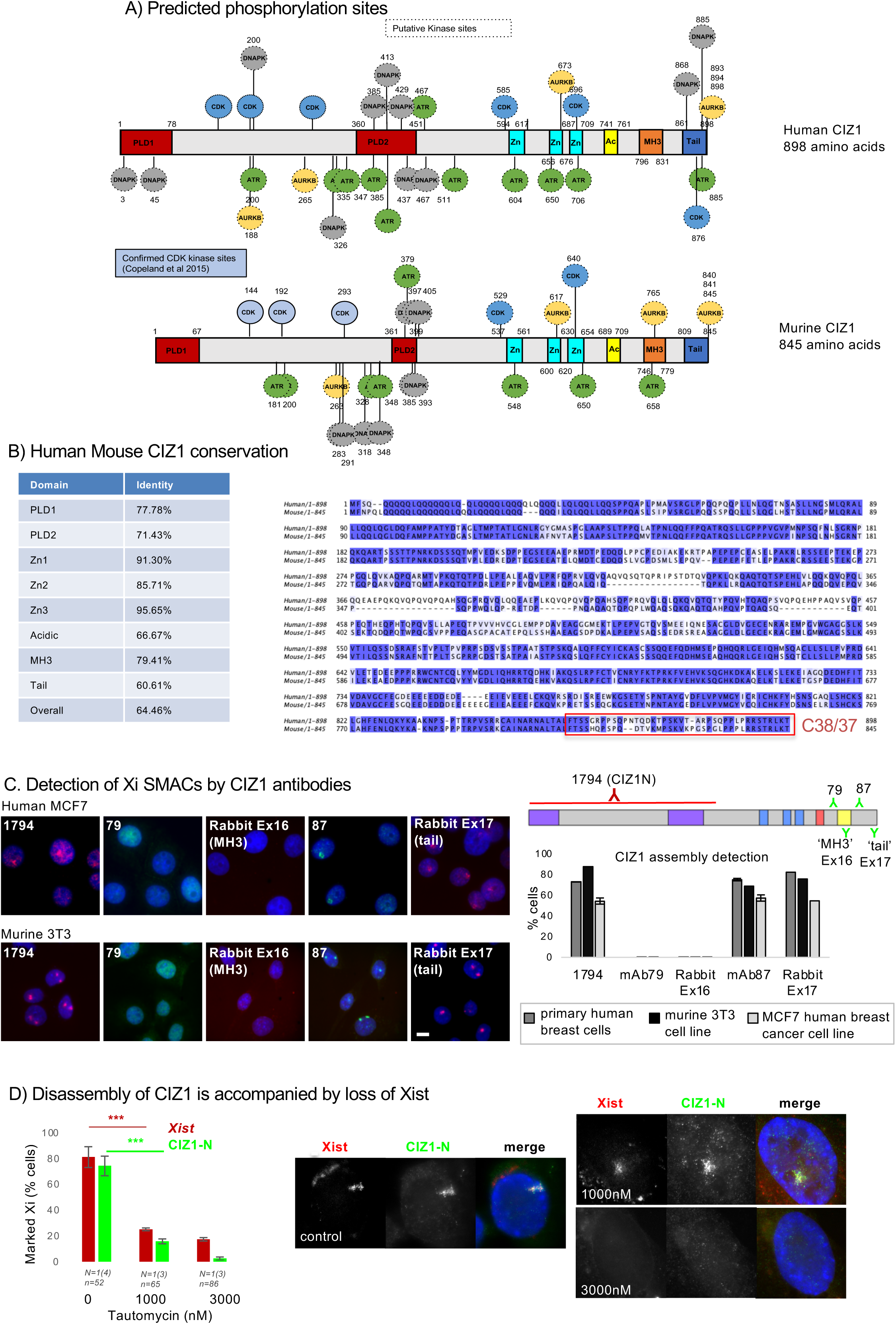
Conservation, functional sites, epitope availability and *Xist* release. A) Putative phosphorylation sites displayed on full length human (Uniprot ID: Q9ULV3-1) and murine (Uniprot ID: Q8VEH2) CIZ1. Data derived from Group-based prediction System (GPS) 5.0 software (18). Consensus AURKB sites (19) are shown in yellow. Functionally relevant CDK phosphorylation sites that were experimentally determined in murine CIZ1 are shown in lighter blue (64). B) Amino-acid alignment, right, showing high primary sequence identity between human and murine CIZ1, with domains in the table, left, created using Jalview 2.11.3.2 software with Tcoffee presets. The C-terminal ‘tail’ 38/37 amino acids are boxed in red, showing that the C-terminal 8 amino acids bearing putative AURKB phosphorylation sites are fully conserved. C) Right, histogram shows frequency of CIZ1 assemblies at Xi in cycling cell populations, comparing five anti-CIZ1 antibodies by immunofluorescence. Above, summary of location of epitopes in relation to CIZ1 domains. Left, example immunofluorescence images showing epitope availability of CIZ1 in Xi assemblies in murine (3T3), and female human (MCF7) cells. D) RNA Immuno-FISH for CIZ1 N and *Xist,* with and without Tautomycin for 15 hours. Histogram shows the proportion of cells in a cycling population that retain CIZ1 or *Xist* at Xi (including tight and dispersed clouds). Right, representative images showing *Xist* (red) and CIZ1-N (green) in D3T3 cells. For 1uM Tautomycin an example of a partially dispersed *Xist* cloud is shown.

**SFig.2.**
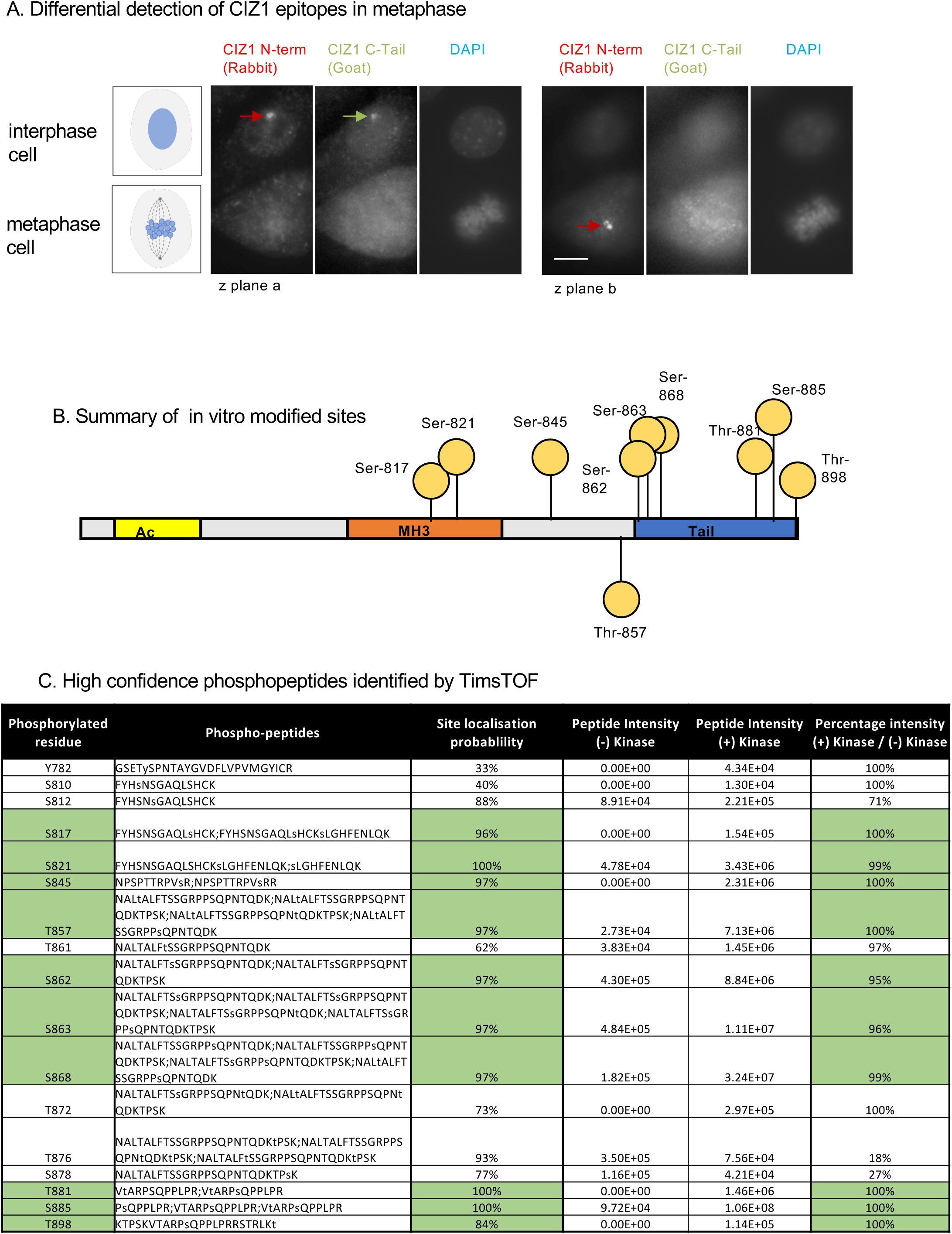
AURKB site epitope availability, and in vitro modified epitopes. A) Example images of neighbouring interphase and metaphase D3T3 cells co-stained for CIZ1-C using a different anti-peptide antibody raised against the tail region encompassing the AURKB kinase site cluster (E17 raised in goat), and CIZ1-N (1794). Two focal planes are shown, revealing both epitopes in the interphase cell (left) but only the CIZ1-N epitope in the metaphase cell. DNA is stained with Dapi. Bar is 10 microns. B) Summary of *in vitro* modified phosphorylation sites showing ten determined sites detected in hCIZ1C179 treated with purified AURKB. C) Table shows identified sites reported from phosphopeptides with ≥95% relative abundance compared to untreated control, and ≥75% site-localisation probability. The phosphopeptides for positive sites/residues are reported in format of lower-case, where t or s is phosphorylated. Residues in reference to full-length human CIZ1 Q9ULV3.

**SFig.3.**
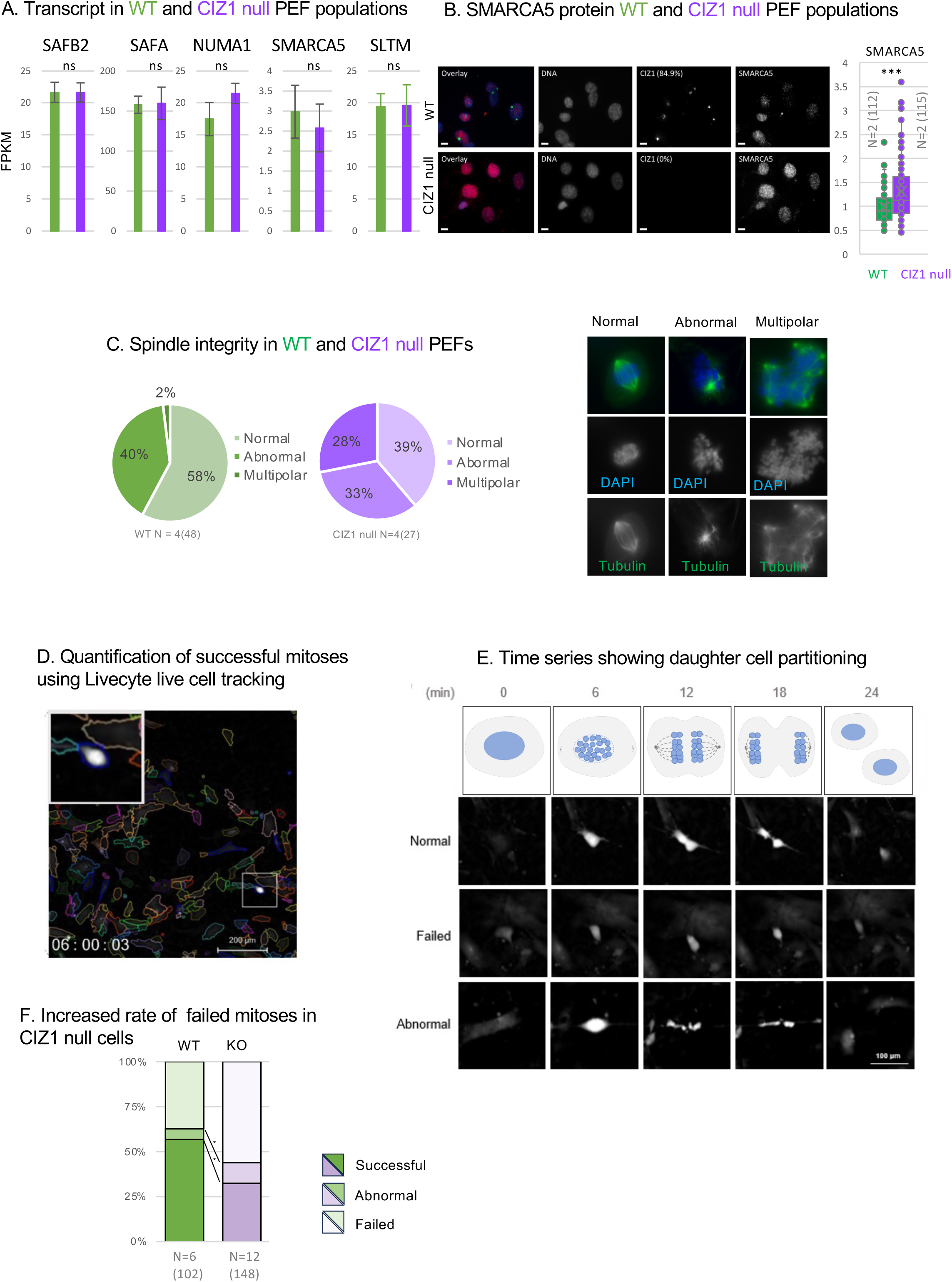
Nuclear matrix proteins and mitotic defects in CIZ1 KO cells. A) Relative transcript levels in triplicate populations of WT and CIZ1 null PEFs (8), showing no significant reduction in expression. B) Example immunofluorescence images of SMARCA5 (red) in cycling WT and CIZ1 null PEFs, co-stained for CIZ1-C (green), and Dapi (blue). Box and whisker plot shows intensity measures derived from two independent primary cell populations (N) for each genotype. n=number of nuclei measured. Comparison is by t-test where *** denotes p<0.001, and indicates a significant elevation of SMARCA5 epitope in CIZ1 null cells. C) Classification of mitotic cells into three groups: cells with visually normal bipolar mitotic spindles, cellsl with abnormal bipolar spindles or aberrant position, or cells with multipolar spindles. Data is expressed as % for mitotic cells derived from four WT and four CIZ1 null asynchronous PEF populations. Below, images show examples of cells stained for alpha-tubulin (ab7291, green), with normal and abnormal spindles. D) Example frame from Livecyte image analysis, monitoring morphological differences and mitotic events in populations of WT and CIZ1 null PEFs, with inset showing a cell undergoing mitosis. Three viewpoints were selected per well for image acquisition over 24 hours. E) Left, example images showing mitotic events that were either normal (giving rise to two daughter cells), abnormal (uneven products), or failed (no separation), seen via timelapse. Right, histogram frequency of each class. CIZ1 null cells have significantly increased rate of failed mitoses, where * denotes p<0.05 (t-test).

**SFig.4.**
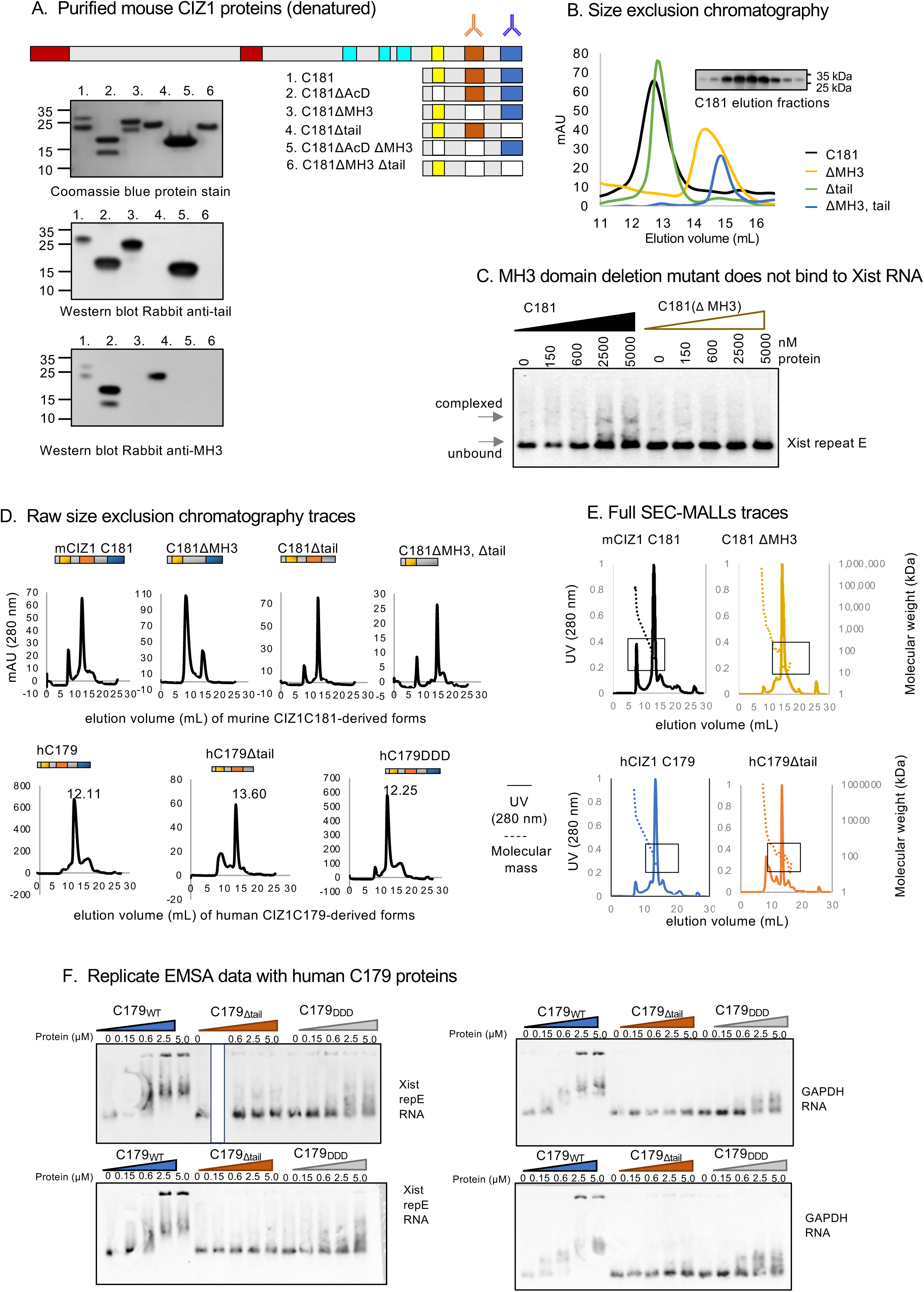
Murine CIZ1 C181-derived proteins, additional EMSA and SEC analysis. A) Murine C181-derived expression constructs. Left, expressed and purified proteins shown after denaturation in SDS-PAGE. Upper gel is stained with coomassie blue to reveal all protein isoforms, and shows the tendency of constructs bearing the 38 amino acid tail domain to produce two forms. Centre, anti-tail antibody fails to detect the smaller of the two in western blot, suggesting that the tail domain is cleaved during expression. Lower, anti-MH3 domain antibody fails to detect MH3 domain deletion mutants, but does detect cleaved and uncleaved forms in which MH3 domain is present. B) SEC profiles for the indicated murine proteins. Inset shows western blot of elution fractions for C181, probed with anti-tail antibody. C) EMSA showing interaction between murine CIZ1-C181_WT_ protein and *Xist* repeat E RNA probe, compared to that with murine CIZ1-C181_ΔMH3_. D) Upper, full SEC chromatograms for purifiied mCIZ1-C181_WT_, mCIZ1-C181_ΔMH3_, mCIZ1-C181_Δtail_ and mCIZ1-C181_ΔMH3Δtail_. Lower, Full SEC chromatograms for purification of hCIZ1-C179_WT_, hCIZ1-C179_Δtail_, and hCIZ1-C179_DDD_. E) Upper, full SEC-MALLS chromatograms showing normalised UV absorbance at 280nm, and molar mass (dotted line) for mCIZ1-C181_WT_, _and_ mCIZ1-C181_ΔMH3._ Lower, full SEC-MALLS chromatograms for of hCIZ1-C179_WT_, and hCIZ1-C179_Δtail_. Boxes indicating where focused versions of graphs are displayed in Fig 6. F) Replicate EMSAs showing *Xist* repeat E (left) and GAPDH (right) RNA interaction with hCIZ1-C179_WT_, and impaired interaction with hCIZ1-C179_Δtail_ or hCIZ1-C179_DDD_. Data, incorporated into quantification shown in Fig.6F.

**Supplemental dataset 1** (Xl). Related to Figures 4 and 5.

Results of three protein interaction studies.

Tab 1 Raw results study A

Tab 2 Raw results study B

Tab 3 Raw results study C

**Supplemental dataset 2.** Related to Figures 4 and 5.

Analysis of three protein interaction studies.

Tab 1 Core 56 interacting proteins

Tab 2 GSEA cluster analysis in STRING, reporting the three highest ranked GO Process and GO Functions.

Tab 3 *Xist* interaction overlap

Tab 4 WT compared to tail deletion mutant

Tab 5 WT compared to phosphomimetic mutant

